# A new field instrument for leaf volatiles reveals an unexpected vertical profile of isoprenoid emission capacities in a tropical forest

**DOI:** 10.1101/2021.02.15.431157

**Authors:** Tyeen C. Taylor, Wit T. Wisniewski, Eliane G. Alves, Raimundo C. de Oliveira, Scott R. Saleska

## Abstract

Both plant physiology and atmospheric chemistry are substantially altered by the emission of volatile isoprenoids (VI), such as isoprene and monoterpenes, from plant leaves. Yet, since gaining scientific attention in the 1950’s, empirical research on leaf VI has been largely confined to laboratory experiments and atmospheric observations. Here, we introduce a new field instrument designed to bridge the scales from leaf to atmosphere, by enabling precision VI detection in real time from plants in their natural ecological setting. With a field campaign in the Brazilian Amazon, we reveal an unexpected distribution of leaf emission capacities (EC) across the vertical axis of the forest canopy, with EC peaking in the mid-canopy instead of the sun-exposed canopy surface, and high emissions occurring in understory specialist species. Compared to the simple interpretation that VI protect leaves from heat stress at the hot canopy surface, our results encourage a more nuanced view of the adaptive role of VI in plants. We infer that forest emissions to the atmosphere depend on the dynamic microenvironments imposed by canopy structure, and not simply on canopy surface conditions. We provide a new emissions inventory from 51 tropical tree species, revealing moderate consistency in EC within taxonomic groups. Our self-contained, portable instrument provides real-time detection and live measurement feedback with precision and detection limits better than 0.5 nmol m^-2^_leaf_ s^-1^. We call the instrument ‘PORCO’ based on the gas detection method: photoionization of organic compounds. We provide a thorough validation of PORCO and demonstrate its capacity to detect ecologically driven variation in leaf emission rates and thus accelerate a nascent field of science: the ecology and ecophysiology of plant volatiles.

**Type of paper:** Method

## Introduction

Forest-atmosphere interactions are shaped not only by the exchange of carbon, water, and energy, but also by the biological production of organic trace gases (Laothawornkitkul et al., 2009; Rieksta et al., 2020; Tani & Mochizuki, 2021; Unger, 2014). For example, volatile isoprenoids (VI)—including 5C isoprene and a diversity of 10C monoterpenes— protect plants from diverse abiotic and biotic stresses (Fineschi et al., 2013), and contribute to the formation of atmospheric aerosols (Heald et al., 2008). These ‘secondary organic aerosols’ formed through atmospheric chemistry with organic gases—most conspicuous in the blue haze that forms over maritime forests in hot conditions—affect the radiative forcing of the climate (Carslaw et al., 2013) and are the source of up to half of all cloud condensation nuclei (Carslaw et al., 2010).

Understanding which plant species emit VI and how they are distributed across landscapes is critical to determining regional and global emissions (P. Harley et al., 2004; Klinger et al., 1998; Purser, Drewer, et al., 2020; Rinnan et al., 2020; Seco et al., 2020). The most abundant organic gas emitted from the biosphere, isoprene, is produced by only approximately one-fifth of terrestrial plant species (Fini et al., 2017), including one-third or more of tropical trees (Taylor et al., 2018). Monoterpene emissions are less abundant but more reactive than isoprene (Heald et al., 2008), and their distribution among species—conspicuous from conifers (pine scent) but cryptic from broadleaf trees—is poorly known (Feng et al., 2019; K. J. Jardine, Zorzanelli, Gimenez, Oliveira Piva, et al., 2020). Models of VI emissions rely on inventories of species measurements from the field to estimate regional fractions of emitting vegetation (Alex Guenther, 2013). This presents a challenge for modeling areas that are under-sampled due to remoteness or high species diversity.

Tropical forests are estimated to emit more VI to the atmosphere, globally, than temperate and boreal forests combined (Alex Guenther, 2013; Hantson et al., 2017; Yáñez-serrano et al., 2020). Variation in species compositions among sites produces substantial variation in emitter fractions, which has been linked to variation in measured forest emissions (P. Harley et al., 2004; Klinger et al., 1998). Given that only one percent of the roughly fifty-thousand species of trees in tropical forests (Slik et al., 2015) have been sampled (Taylor et al., 2018), accurately representing emitter fractions seems a daunting task. The task may be simplified by seeking ecological mechanisms that distribute VI emitting trees across tropical landscapes. For example, a recent study estimated that the proportion of isoprene emitting trees in tropical forests increases two-fold from cool, dry sites to warm, wet sites (Taylor et al., 2018). This pattern may arise due to better photosynthetic performance at high temperatures among isoprene emitting species compared to non-emitting species (Taylor et al., 2019), consistent with the long-held hypothesis that isoprene enhances plant thermal tolerance (Behnke et al., 2007; Sharkey & Monson, 2017; Sharkey & Yeh, 2001; Singsaas et al., 1997). While a promising start, the ecological analyses by Taylor et al. (2018, 2019) relied on published species emissions inventories, most of which were carried out without the aim of ecological hypothesis testing. To both narrow uncertainty in estimates of forest emitter fractions (Taylor et al., 2018), and understand how VI may shape future diversity and function in tropical forests, many more species measurements are needed in contexts that inform the ecological and evolutionary structure of emissions (Russell K. Monson et al., 2013; Sharkey & Monson, 2014).

Despite intensive scientific interest beginning in the 1950’s (Rasmussen & Went, 1964; Sanadze, 2004), research on VI emissions has remained largely confined to laboratory and greenhouse experiments on plants (Sharkey & Monson, 2017), and atmospheric observations and modeling (Sharkey & Monson, 2014). The main impediment to the incorporation of VI into ecological research has arguably been a lack of instrumentation optimized for emission measurements in field settings. There are currently two general approaches to field sampling of leaf VI emissions. The first is an ‘offline’ approach, in which gas samples are collected in the field, and later analyzed in the lab. Typically, leaves are placed in flow-through chambers such as commercial photosynthesis cuvettes or less quantitative enclosures, and gas samples are collected from the cuvette exhaust in bags or adsorption cartridges for later analysis by mass spectrometry (Alves et al., 2014; K. J. Jardine, Zorzanelli, Gimenez, Robles, et al., 2020; Ülo Niinemets et al., 2010; Rieksta et al., 2020). The advantages of this method include control of the leaf environment using a sophisticated leaf cuvette, precision quantification of emissions, and the ability to distinguish VI species. The disadvantages include limited sample sizes due to the high cost of adsorption cartridges, the requirement for offsite analysis, a lack of live measurement feedback, and the potential chemical degradation of stored samples during transport. The second common method employs portable, ‘online’ detection instruments such as handheld photoionization detectors (PIDs), which quantify organic gas concentrations in real time, but lack the ability to distinguish gas species. A portable online detector provides significant advantages in remote areas, affording unlimited sample sizes and live measurement feedback (P. Harley et al., 2004; Klinger et al., 1998). However, low detection precision necessitates high sample concentrations, achieved by high-volume cuvettes that lack environmental control, often enclosing whole branches. Results are typically treated as qualitative, distinguishing only strong emitters from non-emitters.

Here, we present a prototype instrument for precision online detection of leaf VI emissions in the field. We field-validate the instrument with a measurement campaign in the Brazilian Amazon, from which we produce an emissions inventory of 51 tropical trees species, and estimate the vertical distribution of emission capacities within the forest canopy. We call the instrument “PORCO”, after the detection method— photoionization of organic compounds. Custom, light-controlled leaf cuvettes are combined with optimized photoionization detection for high measurement precision and repeatability. The components and methods are adaptable to a range of real-time gas-sampling objectives where required detection limits are around 5 ppb or greater (e.g., greenhouse air, headspace analysis), and to systems requiring microenvironmental control (e.g., moss growth chambers). The adaptations unique to PORCO enable greater sample sizes, lower detection limits, real-time data, and better distinction between leaves and species for more nuanced ecological hypothesis testing than was previously possible.

## Materials and Equipment

### 1 Overview of the ‘PORCO’ system

PORCO measures emissions from intact leaves attached to branches (Fig. 1) in a manner similar to field measurements for leaf photosynthesis (Hunt, 2003). Leaves are enclosed in custom acrylic cuvettes with custom light-emitting-diode (LED) panels providing photosynthetically active radiation (PAR: 90% red, 660 nm; 10% blue, 460 nm) to the leaf to drive photosynthesis and organic gas emissions. Air is purified and pumped to the cuvette inlet at a controlled rate. Cuvette air is subsampled from an outlet while excess flow is exhausted at a loose seal around the leaf petiole. Sample air is drawn through tubing embedded in ice to remove water vapor, and into a commercial photoionization detector (PID). The PID quantifies hydrocarbon gas concentrations, logs its readings at 1 Hz, and displays live numerical data as it is logged. Emission rates from the leaf are calculated as non-steady-state fluxes. The entire system is battery powered and mounted on an external-frame backpack for field portability. For a list of major system components and sources, see Table S1.

**Figure 1:**
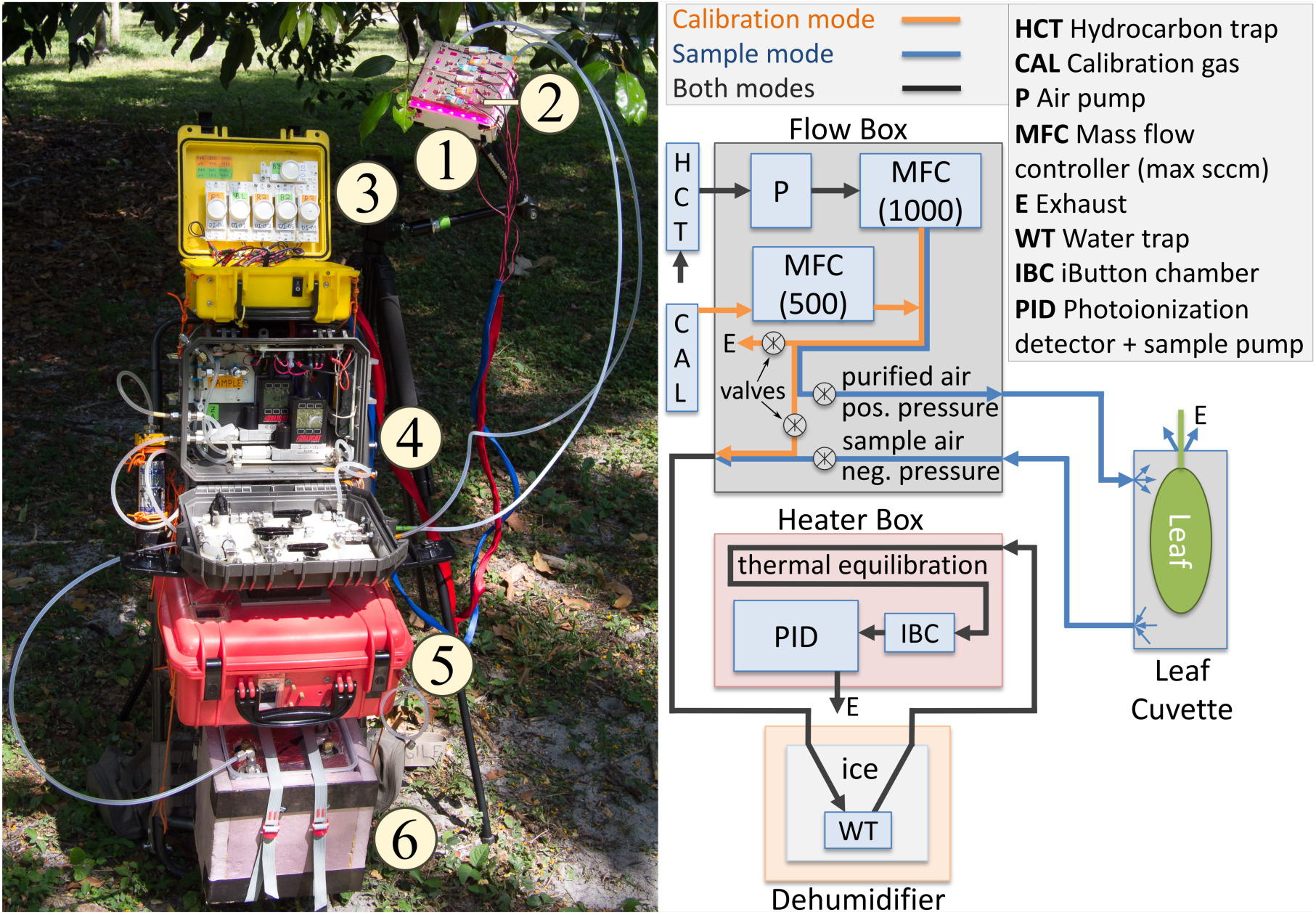
Overview of PORCO components and flow paths. (1) Several acrylic leaf cuvettes accommodate different leaf shapes and sizes. (2) Light emitting diode (LED) panels produce photosynthetically active light. (3) Photon flux density at the leaf is regulated by electrical current control to the light panels. (4) A Flow Box regulates air flow rates and pathways for calibration and sampling via a pump, mass-flow controllers, valve manifold, hydrocarbon trap, and calibration gas. The hydrocarbon trap purifies ambient air for use in calibration and flow-through sampling. (5) Enclosed in a temperature-controlled case, a photoionization detector (PID) with an internal air pump draws sample air from the cuvette. (6) Humidity of calibration air or sample air returning from the leaf cuvette is kept constant by conditioning through chilled tubing in ice and water. The heater, Flow Box, and light control system are each powered by independent lithium ion batteries. The PID contains its own lithium ion battery that powers its internal pump, computer, and sensor components. The PID displays live measurement data, viewable through the enclosure window, in the form of an uncalibrated numerical reading that varies at 1 Hz as it is recorded to the datalog. Data post-processing is performed by custom software in R.

## Methods

### 2 Detecting leaf volatiles by photoionization

PORCO employs a commercial, handheld PID with sensitivity to hydrocarbon gases in parts-per-billion (ppb), the ppbRAE-3000 (Honeywell International Inc., Charlotte, NC, USA). The PID method employs an ultraviolet (UV) lamp to ionize hydrocarbons in the sample gas stream (RAE Systems Inc., 2013c). The separated electrons and positive ions migrate to electrodes in the sensor housing, inducing an electrical current. Gas concentrations are linearly proportional to the induced current.

PIDs have previously been used for qualitative measurements of leaf hydrocarbon emissions in remote field settings, where portability and online detection are significant advantages (P. Harley et al., 2004; Klinger et al., 1998, 2002). The use of PIDs for quantitative emission measurements has been impeded by several factors. Until recently, portable commercial PIDs were limited to parts-per-million (ppm) sensitivity (RAE Systems Inc., 2013c). Measurements suffer from high signal variability (measurement error), often referred to as ‘PID drift’, even when measuring a constant calibration gas (RAE Systems Inc., 2013b). Environmentally controlled cuvettes in photosynthesis machines insufficiently amplify emission signals for detection by PIDs, especially if sources of measurement error are not controlled. The following sections describe the adaptations unique to PORCO that overcome previous limitations to PID use, enabling quantitative leaf emission measurements with high precision and low detection limits in real-time under field conditions.

#### 2.1 PID sensitivities, control, and calibration

PID accuracy is affected by measurement error arising from internal and environmental causes (RAE Systems Inc., 2013b). Measurement error manifests via variation in the ‘baseline’ signal (calibration intercept, or signal from purified ‘zero’ air) and sensor ‘responsivity’ (calibration slope, or baseline-subtracted response to ionizable gas). Both the baseline signal and responsivity vary in response to temperature, humidity, and time (intrinsic drift). In PORCO, PID signal stability and measurement accuracy are enhanced by custom calibration techniques, temperature and humidity control, and data post-processing.

Because measurement error can be positive or negative, monitoring and correcting it requires a continuous non-zero data stream, ideally a raw voltage signal from the instrument. However, commercial PIDs will only log internally processed data based on onboard calibrations. The solution employed in PORCO is to conduct a ‘false calibration’ of the PID that results in its generation of ‘pseudo-raw data’. Pseudo-raw data is the standard PID datalog in putative units of ppb, but calibrated to be a better representation of the raw electrical signal. This returns control of data interpretation to the user, so that corrections for environmental sensitivities and drift can be applied in data post-processing.

The false calibration to generate pseudo-raw data is conducted by performing an onboard ‘zero’ calibration after intentionally reducing the PID’s baseline signal. This can be achieved by one or a combination of the following: apply a zero-air airstream with high humidity; reduce the temperature of the instrument; or install a UV bulb with a lower strength (bulbs of any given model vary in output, and trials will quickly demonstrate which of two bulbs is stronger). All three methods reduce sensor voltage and thereby the signal that the onboard computer interprets as ‘zero’. Upon return to the configuration used during sampling, purified air will read far above zero, thus enabling the perception of any decrease in baseline signal. Similarly, a false ‘span’ (non-zero concentration) calibration can be employed to reduce the minimum signal interval from 1 ppb to less than 1 ppb. For example, conducting a span calibration set to 1000 ppb while providing 500 ppb of calibration gas will induce the instrument to return signal at 0.5 ppb intervals (essentially increasing the sensor ‘gain’). At some point, however, signal noise at the sensor exceeds the apparent resolution of the signal.

To demonstrate the consequences of not controlling for environmental sensitivities and drift, we performed calibration experiments to quantify the effects of humidity, temperature, and time on the PID baseline signal (calibration intercept) and responsivity (calibration slope) (Figs. 2, 3). In the first experiment, the PID was stabilized at a given temperature in a thermally controlled enclosure, and exposed to hydrocarbon-purified air (‘zero-air’) with a series of distinct relative humidity values. Dilutions of isobutylene calibration gas in zero-air were applied, and the process was repeated at a higher temperature (Fig. 2a, b, d). In the second experiment, the PID was stabilized at three different temperatures for a full day each, at constant relative humidity, and isobutylene dilutions were applied every few hours (Fig. 2c, e, f).

**Figure 2:**
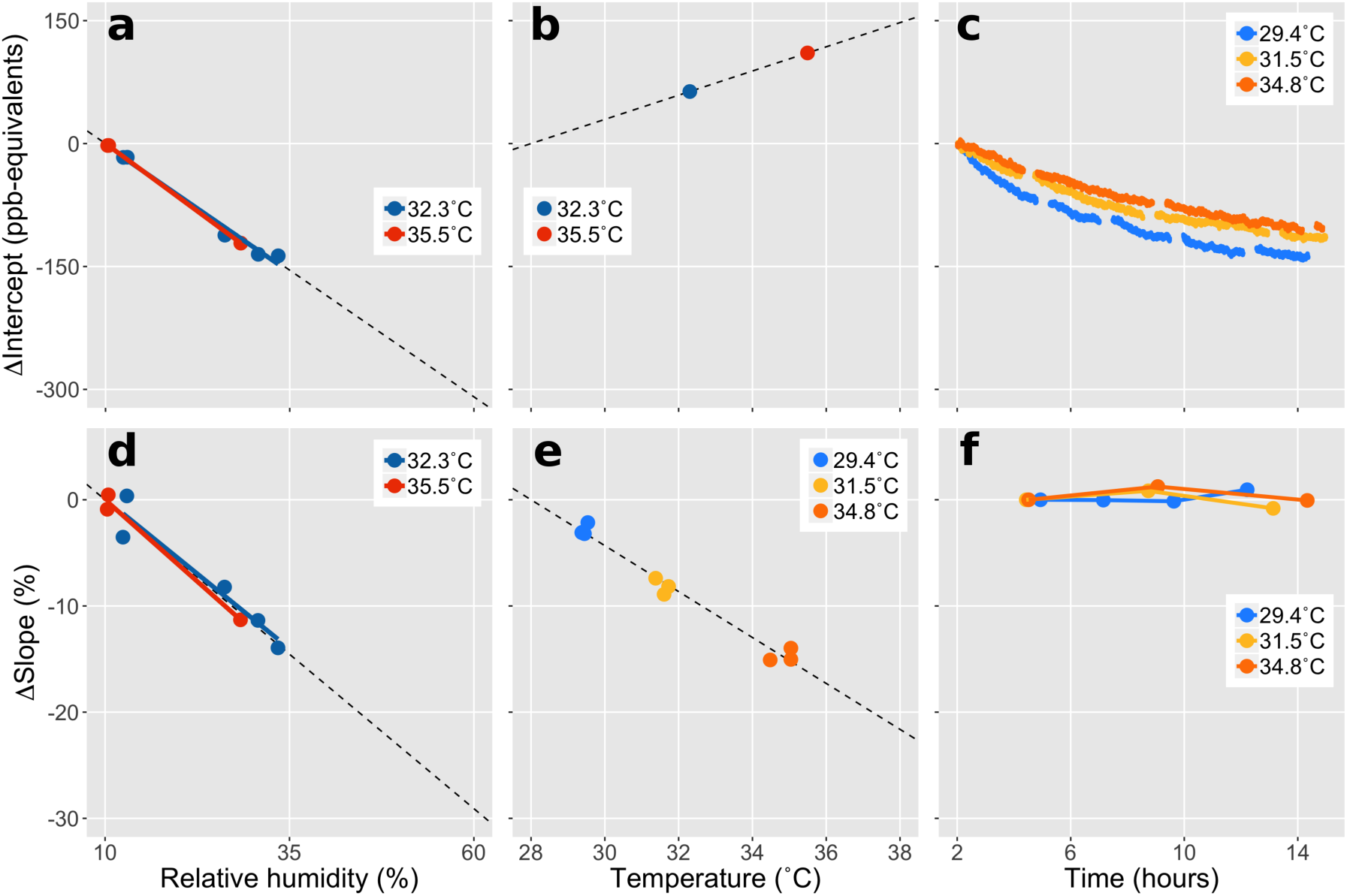
Sensitivity of PID calibration intercept (baseline signal) and slope (responsivity to isobutylene gas) to relative humidity, temperature, and time. To isolate each effect, the results in each panel are expressed as deltas relative to a common reference point. To compare the effect sizes of the different mechanisms of measurement error, the data are plotted across a typical range of diurnal variation of relative humidity (RH, 50 %) and air temperature (T, 10 °C) experienced during outdoor sampling. The negative sensitivity of the calibration intercept to RH (a) is about twice as strong as its positive sensitivity to T (b) and negative sensitivity to operation time (c) over the representative diurnal ranges. The negative sensitivity of the calibration slope to RH (d) is about 1.5 times the negative sensitivity to T (e) over the specified ranges, while no temporal drift was detected (f). We found no interactive effects between T and RH (a, d). To remove the effect of variable RH during T-sensitivity experiments, the calibration coefficients from (b) and (e) were adjusted to represent a common RH based on (a) and (d). Intercepts (a, b) were corrected for temporal drift (c), while the slopes in (e) are derived from three full-day experiments at different temperatures (f).

**Figure 3:**
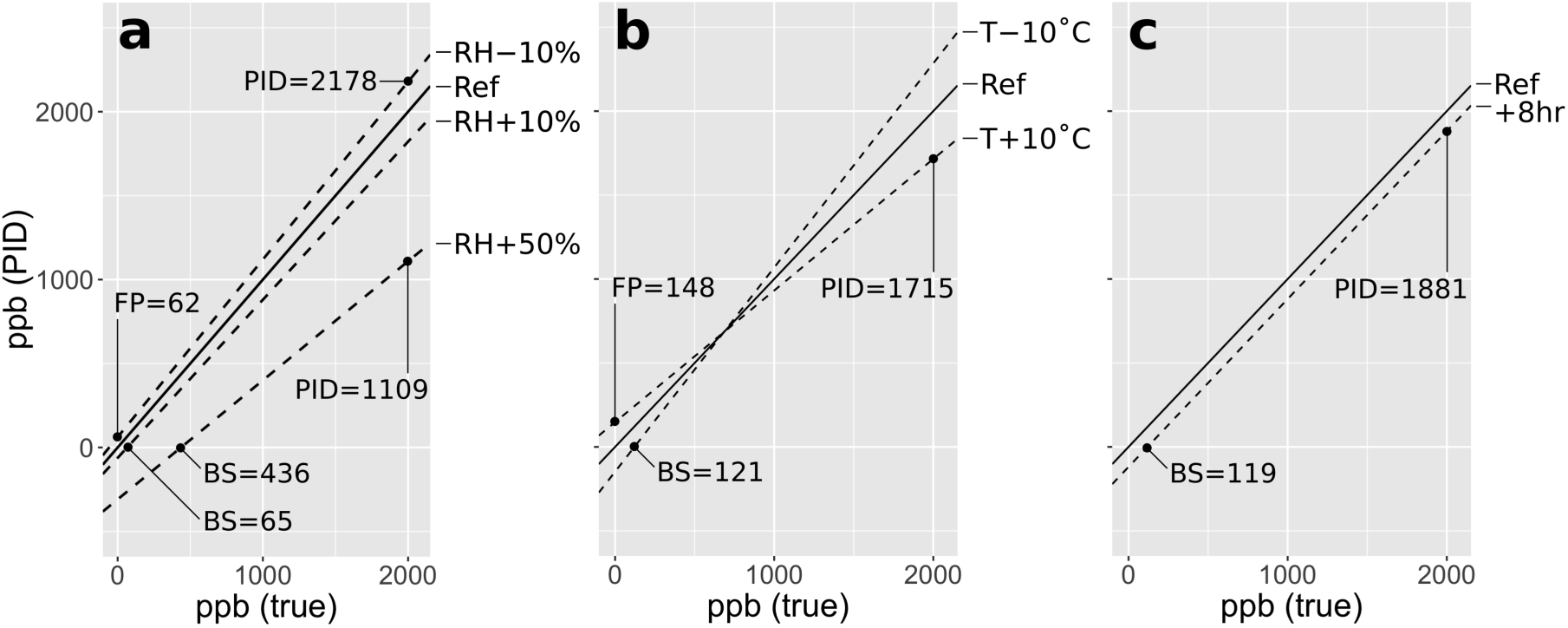
A rough guide for assessing the importance of controlling for PID measurement error (commonly called ‘PID drift’) for a given application and environment. This figure shows estimated outcomes of PID measurements of a gas ranging from 0 to 2000 ppb using the standard onboard calibration method, and measuring under varying sample relative humidity (RH, a), instrument temperature (T, b), and time (c). Conditions are specified at the right of each panel, with ‘Ref’ representing conditions during PID calibration. Sensitivities are derived from Fig. 2. PID measurement error can manifest as ‘blind spots’ (BS) where the PID reads zero for non-zero gas concentrations, false positives (FP) where the PID reads non-zero for zero gas concentrations, and inaccuracies in non-zero gas readings (sensor responsivity). RH will be the most important concern for most users, though direct sun exposure can cause large changes in T. Controlling or correcting for PID measurement error will be most important for applications requiring lower detection limits, precision, and accuracy better than approximately 500-1000 ppb. This should be used only as a rough guide, as PID sensitivities likely differ between sensors.

PID baseline signal was negatively sensitive to relative humidity of the measured air stream (Fig. 2a), positively sensitive to enclosure temperature (Fig. 2b), and negatively sensitive to instrument run time (Fig. 2c). PID responsivity to calibration gas was negatively sensitive to relative humidity (Fig. 2d), negatively sensitive to temperature (Fig. 2e), and not sensitive to instrument run time (Fig. 2f). Extrapolating the modeled sensitivities (linear regressions, dashed lines in Fig. 2) across a 50 % relative humidity range, and a 10 °C temperature range, the relative importance of each factor can be assessed in terms of typical ambient environmental variation encountered during a day of outdoor field sampling. Relative humidity emerges as the most important driver of PID drift, causing a reduction in baseline signal equivalent to 300 ppb (Fig. 2a), and a 30 % reduction in PID responsivity (Fig. 2d) across a 50 % range of relative humidity. This source of measurement error is especially relevant to sampling gases from leaf enclosures, during which transpiration can rapidly increase sample humidity.

The importance of PID measurement error for a given sampling objective depends on the measurement conditions and the lower detection limits and precision required. To facilitate such an evaluation, Fig. 3 shows modeled responses of a PID with a traditional on-board zero and span calibration to gas concentrations between 0 and 2000 ppb. Deviations of PID readings from true concentrations are modeled based on the responses of both baseline signal and responsivity to humidity, temperature, and time shown in Fig. 2. For example, if sample humidity is 50 % greater than calibration humidity, a false zero will be observed up to true concentrations of 436 ppb (the ‘blind spot’), and 2000 ppb will be interpreted as 1109 ppb (Fig. 3a). An instrument temperature 10 °C higher than during calibration will impose a false positive of 148 ppb in purified air, but will underestimate 2000 ppb as 1715 ppb (Fig. 3b). Note especially that the standard calibration technique necessarily imposes a mix of environmental conditions: purified ambient air (at ambient T and RH) is used for the zero, while the span is provided by direct application of calibration gas from the bottle. The calibration gas is both dry (near 0 % RH) and cold due to decompression—the cooling of the sensor causes the typical downward drift in signal observed after completing the span and before disconnecting the PID from the gas bottle. Note that Fig. 3 should be treated only as a rough guide, as sensitivities may vary considerably between sensors and calibration configurations.

While these analyses demonstrate the potential to measure the sources of PID measurement error and apply corrections in data post-processing, the best results are achieved by first maximizing environmental control. For example, temperature measurement of the sample air cannot be used to correct for thermal sensitivity unless sample and instrument are in thermal equilibrium. In PORCO, PID temperature sensitivity is mitigated by controlling instrument and sample-gas temperatures. The PID is enclosed in an insulated case, in which the air is heated to an above-ambient constant temperature (± 0.5 °C) with a thermostat-controlled Peltier device (Fig. 4). Sample (and zero or calibration) gas is equilibrated to the enclosure temperature by routing through coils of stainless steel tubing upstream of the PID (Figs. 1, 4). Relative humidity is held low and constant (approximately 10 % with precision ± 0.5 % at the PID inlet) by routing sample air through tubing embedded in ice and water in an insulated enclosure (Figs. 1, S1), which maintains a saturated air space at constant temperature. This method produces constant humidity over 16 hours of continuous run time in hot (30-40°C air) conditions. A small chamber at the bottom of a V-shaped tubing path in the dehumidifier catches condensed water and allows sample gas to pass above any liquid instead of traveling through it (Fig. S1). Isoprene and other common isoprenoids emitted by plants such as alpha-pinene (a monoterpene) have minimal water-solubility (Kim et al., 2020; Martins et al., 2017), so interference by liquid water is negligible. While some PIDs offer an optional on-board correction for humidity and temperature (toggle in the software; RAE Systems Inc., 2013a), this should be disabled as the correction is small relative to the magnitude of sensitivities, and introduces unaccountable noise into the pseudo-raw data. We find that humidity removal by condensation is the most effective strategy, as our trials with desiccants such as Drierite (W. A. Hammond Drierite Co. Ltd.) showed significant interference with sample gases.

**Figure 4:**
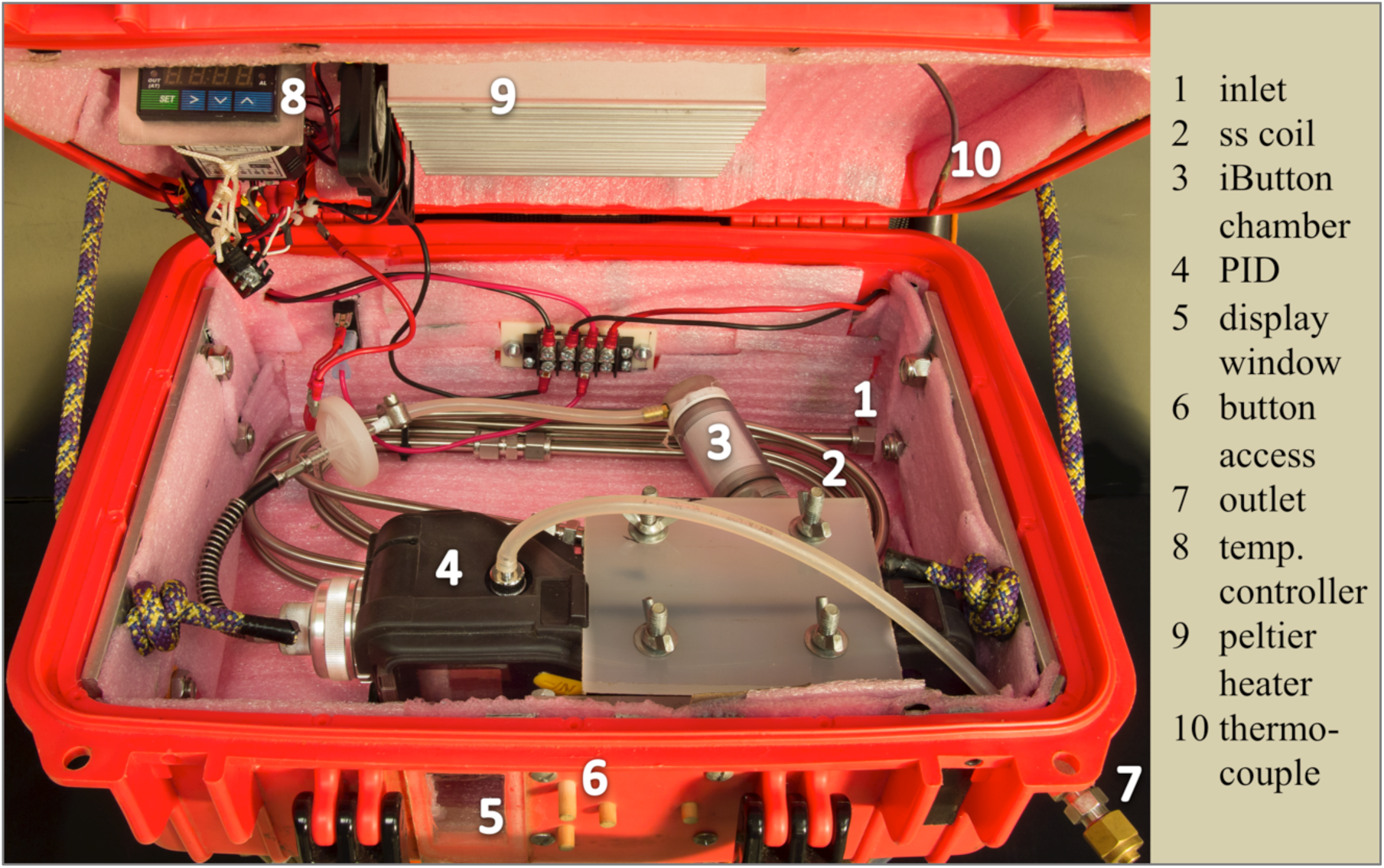
The temperature-control box provides thermal regulation of the PID and sample gas. The PID is fixed inside an insulated enclosure with a battery-powered Peltier heater. A thermocouple-equipped thermostat regulates the heater to maintain a fixed above-ambient temperature inside the enclosure. Sample gas enters the enclosure by tubing connection to a bulkhead union. Sample gas is pre-heated to match the enclosure temperature via coils of stainless steel (ss) tubing. Gas temperature and humidity are measured with an iButton in a PTFE-insulated chamber upstream of the PID. An acrylic window enables viewing of the PID display. Wooden pins at the box interface provide access to the buttons on the PID without opening the enclosure. PID sample exhaust is piped to an outlet port, which can be connected to alternative sampling media (e.g., bags, adsorption cartridges).

PIDs have different responsivities to different gases due to variation in ionization energy among compounds. Measurements of different compounds can be scaled by the relative responsivity of the PID to isobutylene calibration gas using published correction factors (CF) specific to different UV lamp voltages (RAE Systems Inc., 2013a). However, empirical CF may differ between sensors or sample configurations. We find that the PID is 2.25 times more sensitive to isoprene than isobutylene (CF=0.44, Fig. S2), compared to 1.59 times (CF=0.63) reported by Rae Systems (RAE Systems Inc., 2013a). Rarely, some compounds can interfere with PID responsivity (RAE Systems Inc., 2013c). Volatiles from silicone lubricants in valves, for example, can cause chronic signal reduction (‘sensor poisoning’) requiring cleaning of the sensor and sample path to recover responsivity (data not shown). These considerations are particularly important when sample gas contains variable mixtures of compounds, and when choosing materials that contact the sample gas. PORCO employs stainless steel and PTFE or equivalent tubing. Other materials are minimized and tested to ensure non-interference.

#### 2.2 PORCO-PID validation against Fast Isoprene Sensor

We compared the performance of the PORCO-PID to the Fast Isoprene Sensor (FIS, Hills Scientific, Boulder, CO, USA) (A. B. Guenther & Hills, 1998) in a series of tandem calibrations and leaf measurements made in the lab. The FIS uses chemiluminescence in an ozone chamber to isolate the signal of isoprene from other volatiles, and is commonly used for leaf-cuvette based experiments in the lab (Lantz, Solomon, et al., 2019; Russell K. Monson et al., 2016). The instruments were installed in series such that the FIS sampled from the PID exhaust. Isoprene standard dilutions were mixed upstream to generate stepped calibration curves (Fig. 5). Isoprene emission was sampled from four plant leaves, using a LI-6400 leaf cuvette (LI-COR, Inc., Lincoln, NE, USA) for environmental control. The PID sampled from a tee in the cuvette exhaust. Direct calibrations to the FIS showed the inline PID had no effect on FIS calibration linearity or signal-to-noise ratio, but the PID did reduce isoprene concentrations at the FIS by 8.2% (Fig. S3). By calibrating and measuring leaves under the same instrument configuration, signal degradation by the PID was held constant and thereby nullified.

**Figure 5:**
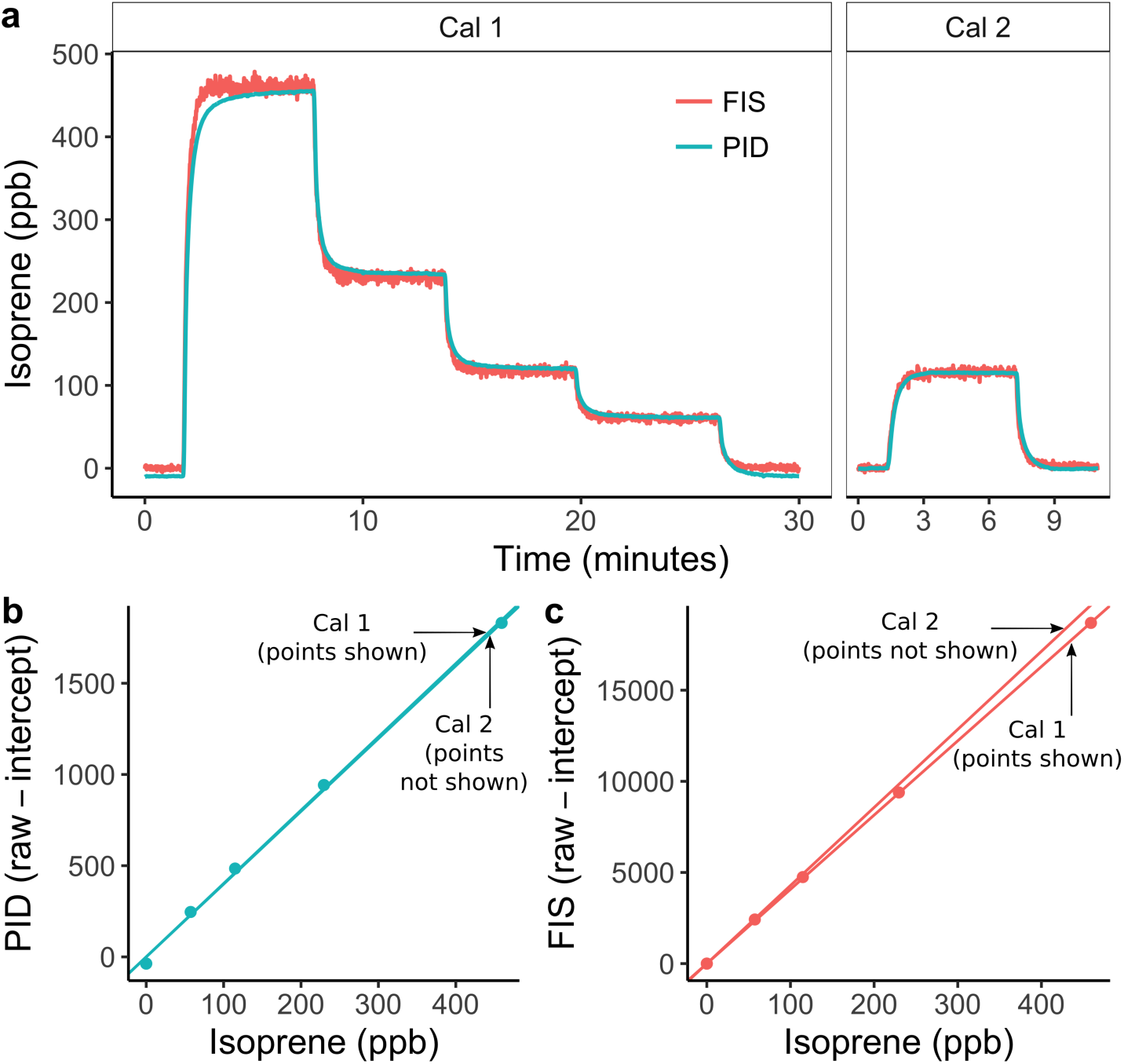
Inter-comparison between the photoionization detector (PID) employed in PORCO and the Fast Isoprene Sensor (FIS) shows comparable precision and signal stability in response to isoprene calibration gas. The PID sampled from an upstream gas mixture, while the FIS sampled from the PID exhaust. (a) Response of each instrument to dilutions of isoprene standard used to produce the calibration curves in (b) and (c). The instrument responses in (a) are visualized in calibrated units of ppb so that instrument signals are aligned and their agreement and stability can be compared. Cal 1 was performed prior to a series of leaf samples, and Cal 2 afterward, with an elapsed time between calibrations of approximately four hours. Points in (a) and (b) are 15 sec and 30 sec averages of PID and FIS raw signals respectively, from the end of each dilution response plateau. Noise at 1 sec resolution is higher in the FIS (a), but the linearity of the FIS response (c) is slightly better than that of the PID (b). PID responsivity (slope) drifted less (b) than FIS responsivity (c), but the FIS background signal (intercept) was more stable over time (not shown, see discussion of temporal PID drift in Main Text).

The PID and FIS showed strong agreement in the time-series of isoprene standard (Fig. 5) and leaf emission measurements (Figs. 6, S4). Compared to the FIS, the PID showed less signal noise at 1 Hz, slightly poorer linearity of response to isoprene standard dilutions, and less drift in responsivity (calibration slope) over time (Fig. 5). The PID showed more drift in baseline signal (not shown, but see Fig. 2c). Both instruments captured the subtle signal of a post-illumination isoprene emission burst (Li & Sharkey, 2013) from an oak leaf (Fig. 6a). Averaged emission rates estimated by the two instruments from the final two minutes of each leaf measurement strongly agreed (linear regression, r^2^ = 0.999, Fig. 7). The PID underestimated emission rates relative to the FIS by 12% (slope [PID∼FIS] = 0.88). The reduction in PID responsivity in sample vs. calibration mode is attributable to a pressure difference between calibration and sample configurations, with a higher flow rate and more resistant excess-flow path producing positive pressure at the PID during calibration (see diagram in Fig. 1 showing positive and negative-pressure flow paths). This discrepancy can be quantified and corrected for any given sample configuration (see *3*.*3 Cuvette Validation*). Best field practices include calibrating and sampling at similar sample-air pressures.

**Figure 6:**
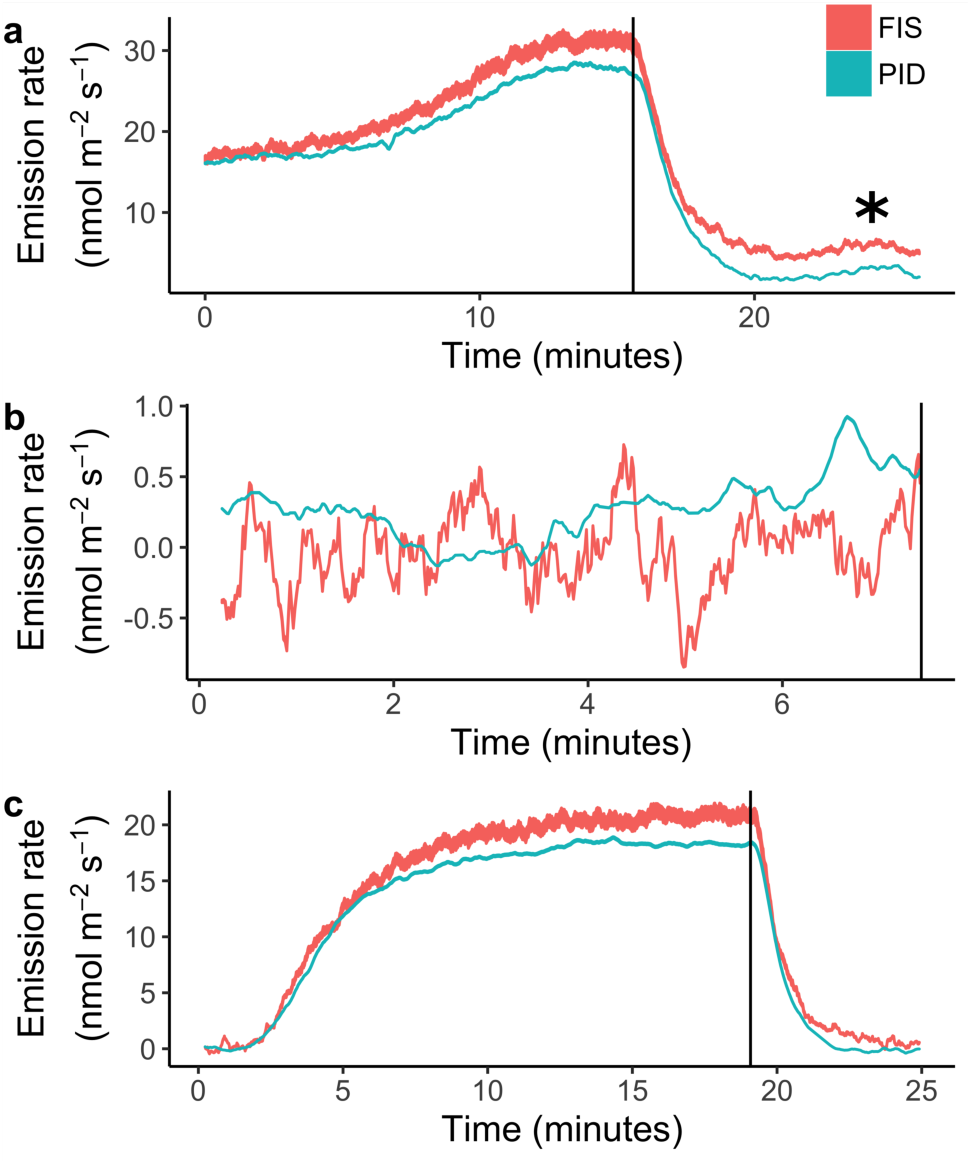
Tandem real-time measurements of isoprene emission from tree leaves by photoionization detector (PID) employed in PORCO, and a Fast Isoprene Sensor (FIS). The PID sampled exhaust gas from a Licor-6400 leaf cuvette, and the FIS sampled from the PID exhaust. Time series data are ribbons, bounded by the emission rates estimated from pre- and post-measurement calibrations, demonstrating confidence relative to sensor responsivity drift. The asterisk in (a) indicates a post-illumination isoprene emission burst captured by both instruments. Species are (a) a temperate oak, *Quercus sp*., (b) an ornamental shrub, *Hibiscus rosa-sinensis*, and (c) a tropical tree, *Malpighia glabra*. Leaf (a) was acclimated to light before emission sampling. For leaves (b) and (c), cuvette light was turned on at time 0. Vertical lines mark the time at lights-off (cuvette was opened at lights-off in (b)).

**Figure 7:**
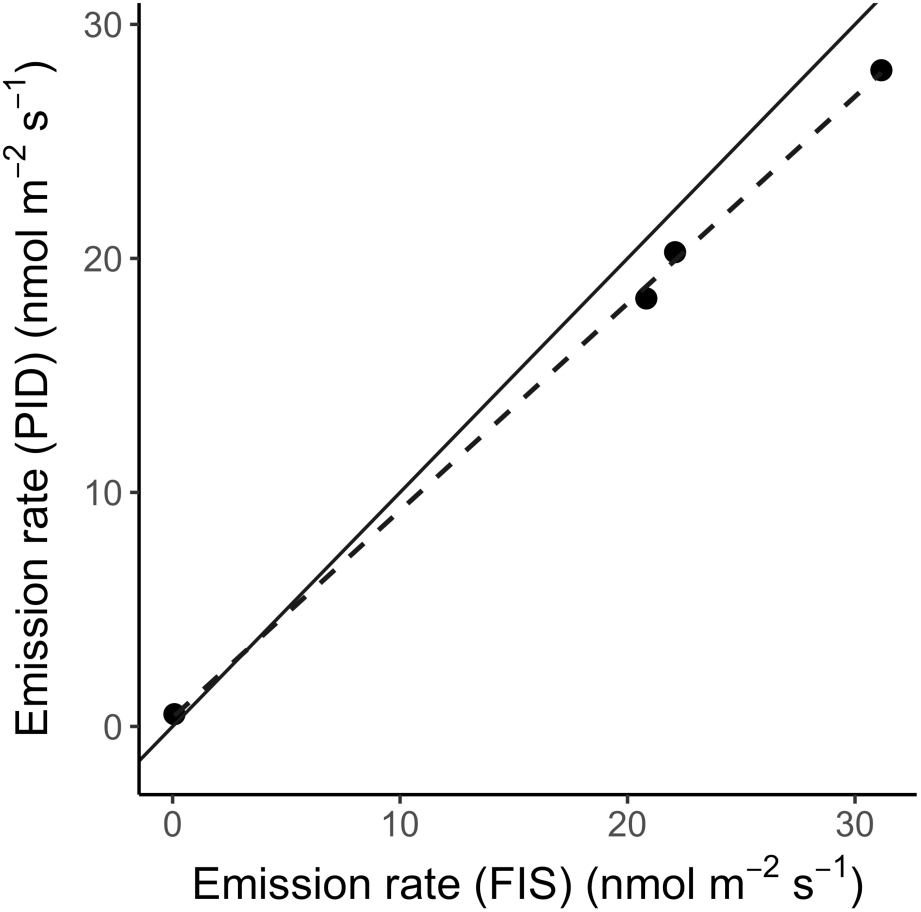
Isoprene emission rates calculated from photoionization detector (PID) and Fast Isoprene Sensor (FIS) readings, averaged over the final two minutes of leaf illumination from four leaf samples (including those depicted in Fig. 6 and an additional sample from *Malpighia glabra*, Fig. S4). The solid line is 1:1. The dashed line shows a linear regression through the data points (r^2^ = 0.999, slope = 0.88).

### 3 Leaf cuvettes

Cuvettes used for capturing volatile emissions from plant leaves vary from translucent bags enclosing whole branches (Padhy & Varshney, 2005; Geron *et al*., 2016) to highly environmentally controlled leaf enclosures such as those employed in commercial photosynthesis instruments (Alves *et al*., 2014; Jardine *et al*., 2015). While PORCO-PID adaptations can improve real-time field sampling precision from a diversity of cuvettes, PORCO employs custom leaf cuvettes with design features that optimize precision and detection limits relative to the sensitivities of the PID. The cuvettes also provide a low-cost solution for quantitative leaf emission sampling using other gas-detection devices. Leaf illumination can be provided from ambient light, or using the custom modular LED panels described in Section *4: Leaf light environment*.

#### 3.1 Cuvette design

To quantify emission rates, sampled trace gas concentrations must exceed the lower detection limit (LDL) of the detector. Emitted gases accumulate inside a cuvette at a rate dependent on the leaf size, emission rate, cuvette volume, and the replacement rate of cuvette air. The PORCO cuvettes were designed to achieve a LDL equal to a leaf area-based emission rate (ER) of 1 nmol m^-2^ s^-1^ or better within 60 sec of measurement from a majority of leaf types found among tropical rainforest trees. A 60 sec LDL of ER = 1 is a conservative goal that allows for confident distinction of weak emissions from non-emission over slightly longer (minutes) measurement times under non-ideal conditions such as sub-optimally sized leaves, reduced physiological activity of cut branches, or environmental instability causing PID measurement error. We evaluated cuvette designs in terms of the ratio of ER to the concentration of the emitted gas accumulated in the cuvette after 60 s, which we term the ‘signal amplification factor’ (SAF). Given typical signal noise of up to ± 2 ppb of isoprene, a confident LDL for the PORCO-PID is approximately 5 ppb. Therefore, a minimum SAF = 5 is optimal for detection of ER = 1 (i.e. 5 ppb is accumulated in 60 s). For comparison, for the LI-6400 leaf cuvette, SAF = 1.5 due to the small cuvette volume relative to a standard flow rate (550 sccm), resulting in LDL = 3.3 nmol m^-2^ s^-1^. Moreover, at a given emission rate, gas concentrations in the LI-6400 cuvette reach steady state in less than 1 min, while concentrations continue to increase in larger cuvettes beyond 1 min, giving a greater potential to exceed lower detection limits.

Because the flow rate through the cuvette must exceed the sampling rate of the PID (300 or 500 mLpm), increasing SAF requires increasing the amount of leaf area enclosed, while minimizing the increase in cuvette volume. We designed three cuvettes that provide SAF > 5 by fully enclosing entire leaves or leaflets (of compound leaves), with distinct shapes that accommodate leaf types of a large majority of tropical tree species.

#### 3.2 Cuvette construction

Custom PORCO leaf cuvettes (Fig. 8) are rigid acrylic boxes that enclose entire leaves. The petiole or twig traverses a port by which leaves maintain connection to their branch. Operation is a flow-through system where pure air is pumped through an inlet port, and sample air is drawn by the pump in the PID via an outlet port (diagram in Fig. 1). Excess flow is exhausted through a partial-seal around the leaf petiole (‘petiole gasket’, Fig. 8). An internal fan ensures even air mixing, necessary for non-steady-state flux calculations. All internal parts are acrylic, stainless steel, or PTFE to minimize adsorption and release of trace gases, except for the plastic mixing fan, which does not produce significant interference. The acrylic initially produced significant background concentrations of ionizable gases, but this was successfully eliminated by baking cuvettes in a drying oven at 60 °C for five days. A roof mounting system accommodates modular LED panels and a light diffusor. Reflective siding ensures even light distribution and minimal interference from ambient light.

**Figure 8:**
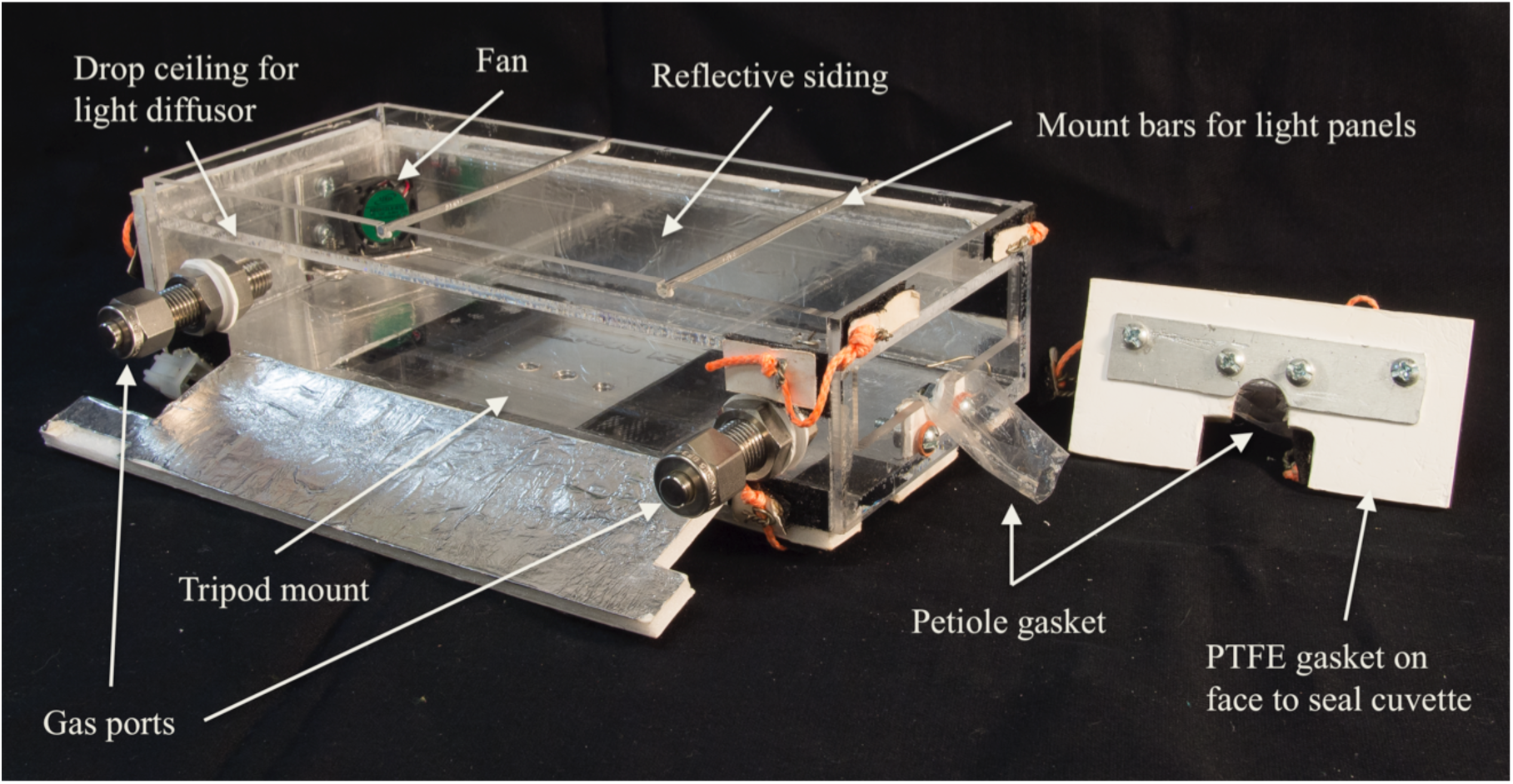
One of three uniquely shaped leaf cuvettes, designed to accommodate a range of leaf shapes and sizes. The leaf is inserted into the opening at right, and the petiole settled into the U-shaped slot. Tensioned cords seal the faceplate with the PTFE gasket to the front of the box, with its inverted U-shaped slot and petiole gasket opposing those on the cuvette face. The petiole gasket is loosely sealed around the petiole with cord such that excess flow (150 mLpm or greater) can escape, but diffusive gas exchange with outside air is not permitted. Light panels mount to the top of the cuvette to drive photosynthesis. Reflective siding ensures even light distribution and minimal interference from ambient light.

#### 3.3 Cuvette validation

Constant leaf emission inside a cuvette should produce an asymptotic increase in gas concentration. The shape of the concentration curve depends on the emission rate, cuvette volume, and flow rate. With a known flow rate and cuvette volume, the emission rate can be calculated from any portion of the curve. Assumptions about the shape of the curve can be validated by simulating leaf emissions by conveying calibration gas to the cuvette inlet (Fig. 9). Insufficient air mixing would reduce the effective mixing volume, producing an initial increase in concentration that is faster than expected. Our simulated emissions show an initial increase that is slightly slower than expected. Insufficient excess flow out of the petiole gasket may produce a diluted curve if the seal is poor, due to mixing with outside ambient air. However, tightening the petiole gasket seal or increasing the flow rate to the cuvette does not alter the results (data not shown), suggesting that excess flow through the loosely sealed petiole gasket under normal sampling conditions is sufficient to mitigate dilution.

**Figure 9:**
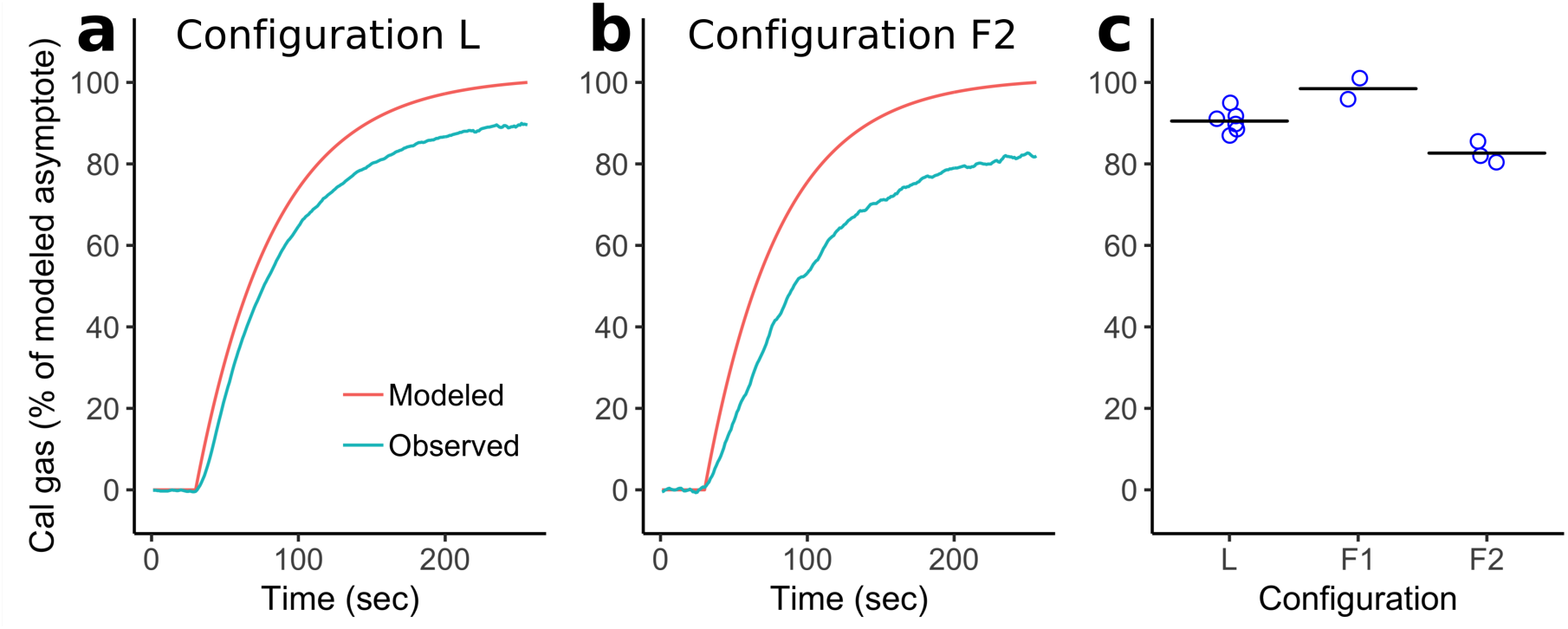
Calibration gas is piped to leaf cuvettes to simulate leaf emissions in order to validate emission calculations from curves of concentration changes. Calibration correction factors are derived from emission simulations to correct for the effect of pressure differential between calibration and sample modes on PID sensor readings. (a) Example of a simulated emission showing expected (modeled) and observed cuvette concentrations measured by PID under a laboratory configuration. (b) Example of a simulated emission under the F2 field configuration. (c) Observed concentrations at 210 sec as a percentage of expected concentrations during simulated emissions under three different plumbing configurations, one lab, two field. Mean values (horizontal lines in (c)) are used to correct sample readings for the effect of the pressure drop for a given sampling configuration.

Simulated emissions show that the largest measurement bias arises from pressure differences caused by different plumbing configurations. The balance of pressure from the zero-air pump (positive) and PID pump (negative) differ between calibration and sample modes (Fig. 1), and between different sampling configurations. During calibration, positive pressure is determined by the total flow rate and the resistance of the excess-flow exhaust path (coiled 1/8” tubing). During sampling, the PID pump pulls air from the cuvette through a relatively long pathway via the dehumidifier and thermal equilibration tubing (Fig. 1), causing a pressure drop. We have not attempted to measure pressure in order to quantify this effect. Pressure effects are particularly evident when changing flow modes by turning valves, which inevitably causes a transient spike in PID readings (Fig. S5). If valves are turned too quickly, or the flow path is momentarily blocked, this reading spike can be extremely large and persist in the form of an asymptotic recovery toward the previous baseline signal over a half hour or more. Strong negative pressure could also exacerbate the effect of small leaks in sample path connections, though we note that sample dilution caused by leaks is inconsistent with the observation of reduced signal at the PID (after switching from calibration to sample mode) relative to an independent instrument sampling directly from the PID exhaust (i.e. the independent sensor should have seen the same reduction in signal, see Section *2*.*2 PORCO-PID validation against Fast Isoprene Sensor*). Pressure effects are plausibly related to the fact that the PID pump, ionization bulb, and sensor are all powered by the same battery, and therefore a change in pump effort caused by a change in flow pressure causes a change in sensor voltage. Different calibration and sample configurations therefore produce a corresponding offset of observation from expectation during simulated emissions. The offset is consistent within configurations (Fig. 9c), allowing calculation of a configuration-specific correction factor for leaf emission estimates. Calibrating exclusively via simulated emissions would obviate the need for correction factors, but this requires much more calibration gas, which is in limited supply in the field due to the logistical challenge of transporting compressed gas cannisters.

### 4 Leaf light environment

PORCO is designed to measure emissions of light-dependent hydrocarbons, i.e. those responsive to light via the dependency of their production on photosynthetic metabolism (Laothawornkitkul et al., 2009). For comparability of measurements, leaf illumination by photosynthetically active radiation (PAR) is typically standardized to 1000 umol photons m^-2^ leaf s^-1^ (R K Monson et al., 1995; Ü Niinemets et al., 2010). In PORCO, PAR is provided by custom panels of light emitting diodes (LEDs) mounted atop the leaf cuvettes (see Section *6*.*1: Standardized leaf measurement with PORCO*). The light panels were designed to mimic the peak light wavelengths and ratios employed in the LI-6400 leaf chamber: 90% red light with peak output at 670 nm, and 10% blue light at 465 nm. The PORCO light panels use Cree Xlamp “Photo Red” (peak output at 650-670 nm) and “Royal Blue” (450-465 nm) LEDs (Cree Inc., Durham, NC, USA). Light ratios and total quantity are regulated based on a calibrated relationship between measured PAR output and electrical current to the bulbs (Fig. S6). During measurements, electrical current to each bulb color on each panel is adjusted to target values via dimmer dials, based on voltage read across precision inline resistors (yellow box in Fig. 1). Electrical current-controllers maintain a steady photon flux density at the leaf plane.

The modular LED panels can be adapted to a diversity of cuvettes. In PORCO, up to three panels are used at a time depending on cuvette size. The LED panels could be easily applied to custom cuvettes adapted for other purposes, such as moss or algal growth chambers.

### 5 Field calibration and data post-processing

The continuous pseudo-raw data logged at 1 Hz from the PID is processed with custom code in R (R Core Team, 2020; Wickham, 2016). Fig. 10 illustrates the processing flow from a day of field measurements. During measurements, the user records start and end times for each type of sample, such as zero-air, calibration air, cuvette blanks, and leaf samples. Data processing is applied to specific time brackets in the time-series output based on the type of measurement recorded.

**Figure 10:**
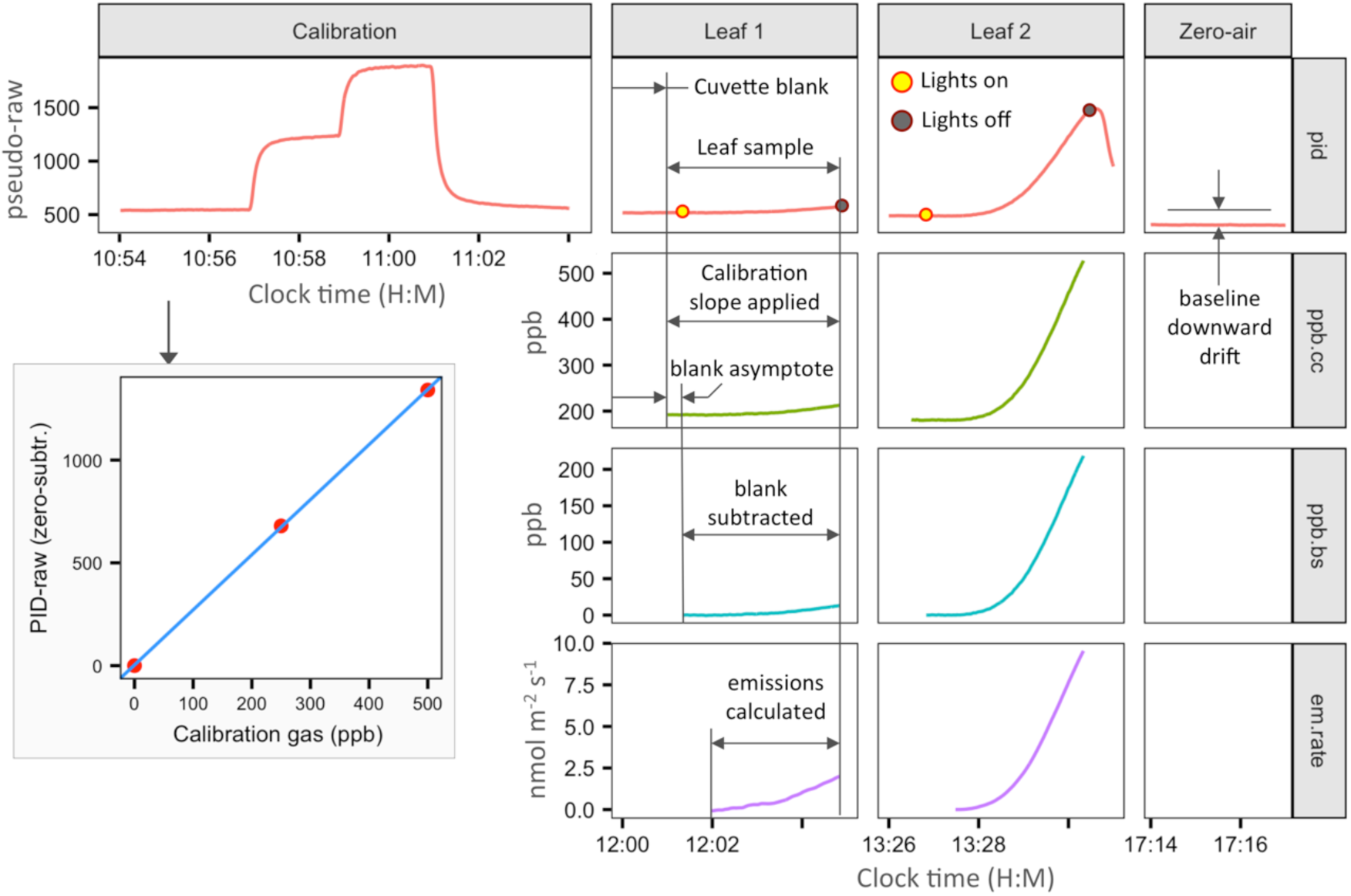
Demonstration of the data processing flow from selected time segments during a day of measurements. Data shown is actual field data from a tropical rainforest. The timing of all calibration and sampling activities are manually noted with 1 sec resolution and entered into standardized tables for data processing. Custom processing code in R applies calculations and data transformations according to the activity specified for each time bracket. A calibration determines the responsivity coefficients for conversion of PID pseudo-raw data to gas concentrations. With a leaf in the cuvette, the background is allowed to stabilize for several minutes. Leaf measurement data is converted to calibration-corrected ppb (ppb.cc) by the calibration slope only. The asymptotic tail of the cuvette blank (stabilization period with the leaf in darkness) is then subtracted from the subsequent leaf-sample period (ppb.bs). The relative responsivity to isoprene (0.44 compared to isobutylene calibration gas) is also applied. Emission rate (em.rate) is then calculated as a non-steady-state flux from the time-series of ppb.bs data.

Linear calibration curves are made by measuring isobutylene standard diluted in purified (‘zero’) air. The air streams are regulated at desired flow rates by mass-based flow controllers (Alicat Scientific, Tucson, AZ, USA) and joined before exhausting through coiled eighth-inch tubing to minimize ambient interference, while the PID samples the mixture from a tee junction. Calibrations are performed at the beginning of each day of measurements, and frequently at the end of each day to ensure responsivity stability. Periodic zero-air readings can be used to interpolate and subtract baseline drift (Fig. 2c).

For standard leaf measurements with PORCO cuvettes, the stabilized signal from the cuvette with the leaf installed and lights off (‘cuvette blank’) is used as the baseline for emission calculations. Leaf emissions are calculated as a non-steady state flux from the rate of change in concentration relative to the blank, determined by linear regressions on a 40 sec sliding window of data following the moment the cuvette lights are turned on (Fig. 10).

### 6 Integrated system

Integrating all of these components into a single, portable system allows us to bring the laboratory to the leaf. For the first time, a system that can be carried and operated by a single person under field conditions can provide precise, real-time information about plant volatile emissions, allowing unlimited sampling. In contrast to ‘offline’ sampling methods requiring subsequent laboratory analysis, PORCO provides live measurement feedback and the ability to view processed results after each measurement day, allowing the user to make informed adjustments to their sampling efforts while in the field. PORCO has the potential to accelerate a nascent field of science: the ecology and ecophysiology of plant volatiles.

#### 6.1 Standardized leaf measurement with PORCO

PORCO measures emissions from leaves that are attached to branches and under conditions suitable for photosynthesis (Fig. 11). Measurement preparation begins with choosing a cuvette for the leaf that maximizes the ratio of leaf area to horizontal cuvette area without bending the leaf edges. Purified ambient air is conveyed to the cuvette inlet port via a mass-flow controller, while the PID draws air from the outlet port. The LED panels are attached to the cuvette, and electrical current is adjusted to produce the desired light output (1000 PAR for standardized measurements). The lights are then switched off.

**Figure 11:**
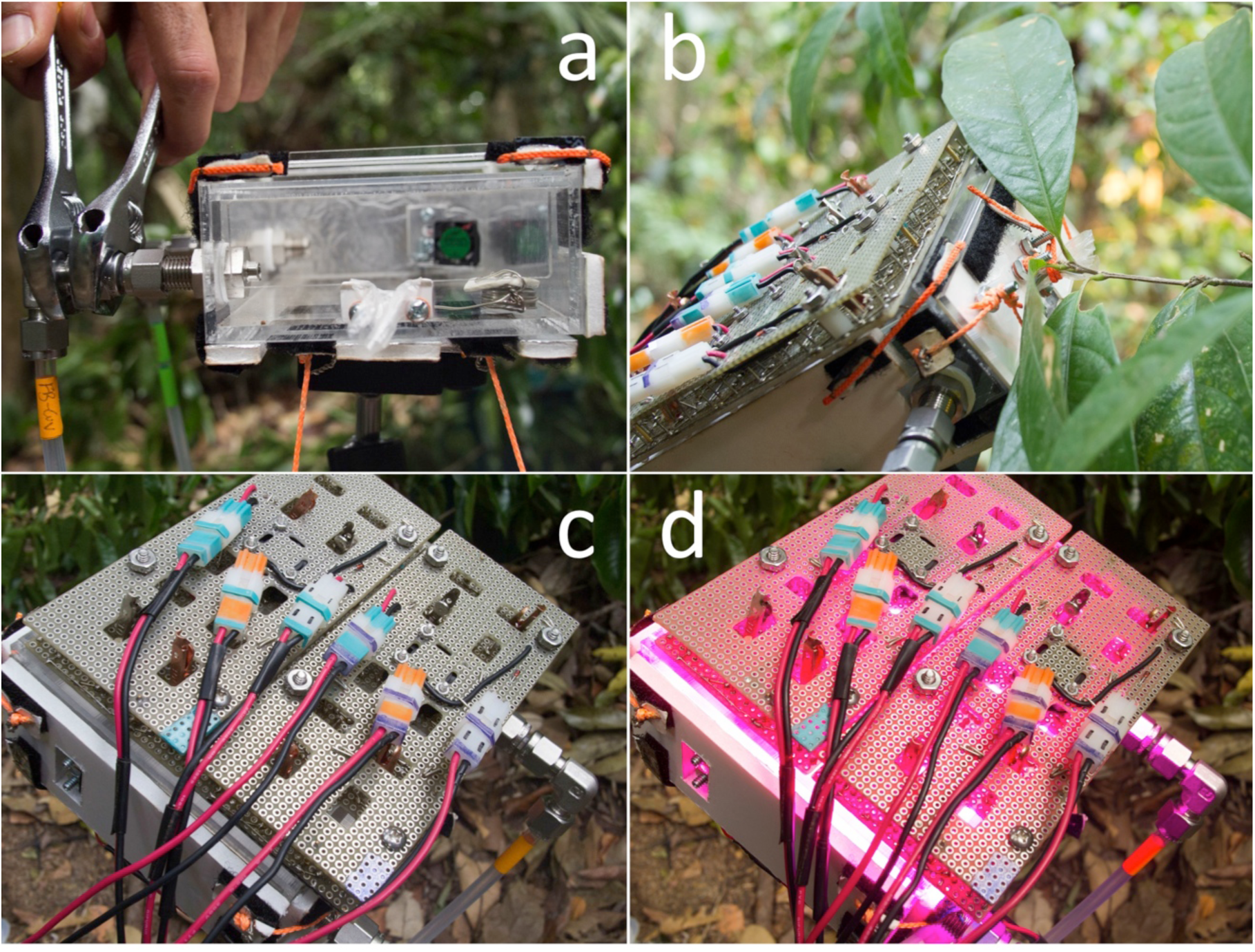
To measure emissions from a leaf, the appropriate cuvette is chosen (a), maximizing the amount of leaf area relative to cuvette volume. The leaf is installed while still attached to its branch, and the cuvette faceplate fixed with the petiole gasket loosely sealed around the leaf petiole (b). The leaf sits in darkness until ambient air is cleared (c). Then the lights are switched on, set to deliver 1000 umol m^-2^ s^-1^ PAR (d).

The leaf is installed by carefully positioning the cuvette to enclose the leaf, using an articulating and locking arm mount to closely match the cuvette orientation to leaf orientation, minimizing leaf disturbance. The cuvette faceplate is installed and the petiole loosely sealed by the petiole gasket. Purified air is conveyed to the cuvette at a rate ≥ 150 sccm above the PID sampling rate (either approximately 500 mLpm or 300 mLpm depending on PID pump-speed setting). Excess flow is exhausted from a loose seal around the leaf petiole to resist diffusion of ambient air into the cuvette. The leaf is left in darkness inside the closed cuvette for sufficient time to clear ambient air and achieve a stable background signal (6-9 min depending on cuvette volume). The LED lights are then switched on, driving photosynthesis and VI emissions if the leaf is an emitter.

For standardized measurements at 1000 PAR, leaves are illuminated for 3.5 min. Longer illumination periods at high light levels are avoided due to the potential to overheat the cuvette. Continued monitoring following lights-off allows confirmation of weak emission peaks, and analysis of species variation in post-illumination emission dynamics (e.g., Fig. 6a). Live emission information can be qualitatively determined during a measurement by monitoring the numerical output on the PID display (a single number varying at 1 Hz, equivalent to the datalog) through the enclosure window (Fig. 4). Careful observation allows the distinction of even weak emitters from non-emitters in real time, as well as the detection of anomalous signal drift that would contribute to a bad measurement. When measurements are completed, leaves are scanned to determine their area, then dried and weighed so that emission rates can be scaled by either leaf area or mass.

#### 6.2 Field measurements from 51 tropical tree species

The primary components of PORCO, excluding leaf cuvettes, are mounted on an external-frame backpack for portability. The system can be carried by one person through a forest, up a canopy tower, or hauled by ropes into trees or canopy walkways (Fig. 12). All of the plastic cases can be closed during transport with all air-flow components still running to maintain PID stability. For tall trees, branches can be cut, carefully lowered, re-cut under water to maintain the transpiration stream, and then measured on the ground. Where accessible, the portability of PORCO allows in-situ measurements of tree leaves, which eliminates the potential for undesired physiological responses to branch cutting.

**Figure 12:**
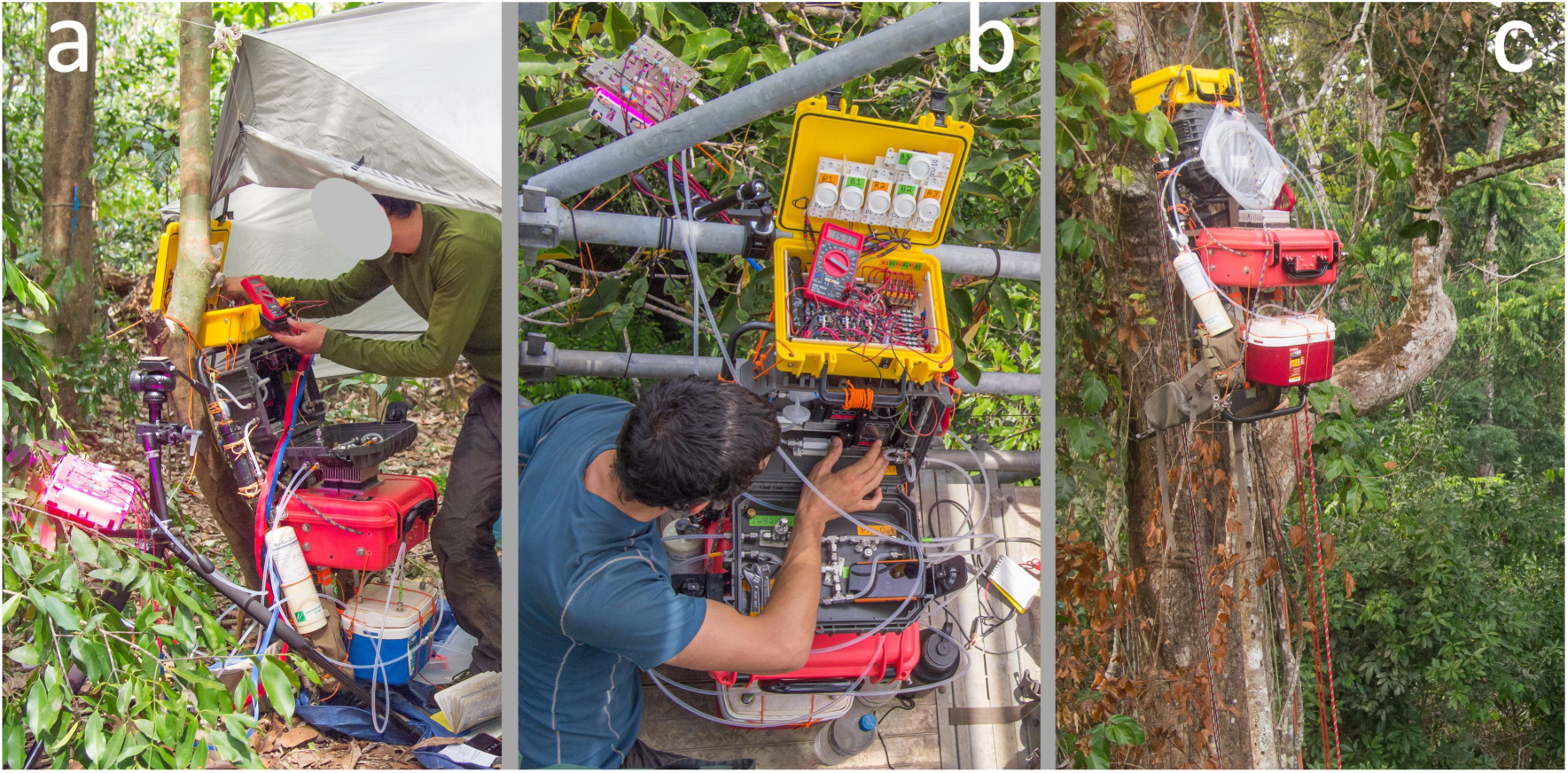
Field measurements with PORCO in a tropical forest: (a) on the ground with cut branches; (b) on a canopy walk-up tower for *in situ* measurements on intact branches; (c) hauled into the canopy for *in situ* measurements from a suspended walkway. (Identifying features of persons in photos has been obscured as per bioRxiv preprint policy.)

We used our uniquely portable detection system to quantify leaf VI emissions from tropical trees at a field site in the eastern Amazon, the “k67” eddy flux tower site in the Tapajos National Forest near Santarém, PA, Brazil (Rice et al., 2004). In 27 days of field sampling, we obtained high quality emission data from 138 leaves from 67 trees and 51 species (Table 1). All species were professionally identified from pressed specimens (Herbario IAN of Embrapa in Belém, PA, Brazil), and taxonomic names were standardized by the Taxonomic Name Resolution Service (Boyle et al., 2013) (accessed Aug. 22, 2020, Table S2).

**Table 1:**
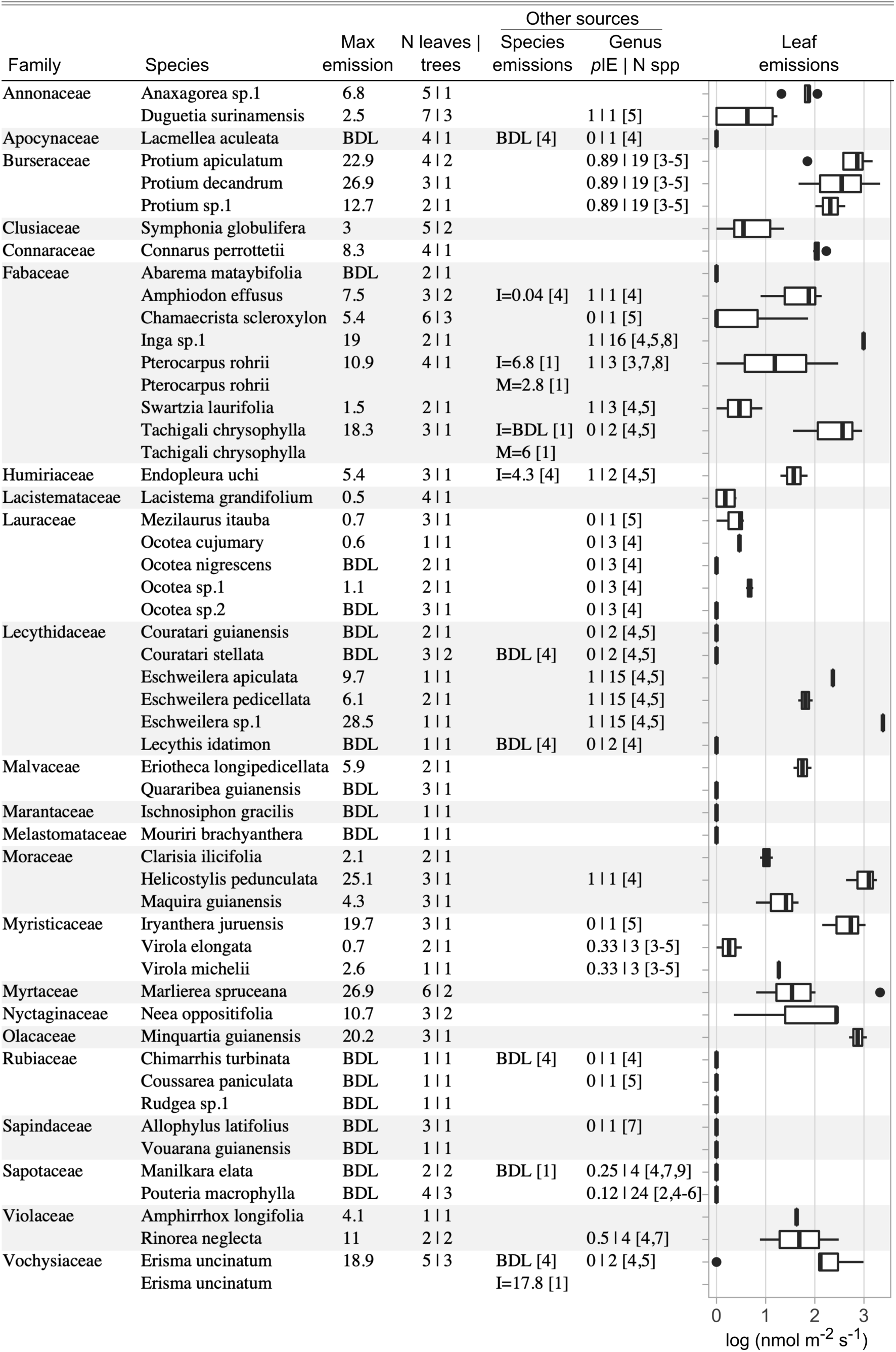
Volatile isoprenoid emission rates from 51 tree species at an eastern Amazon site, the Tapajós National Forest, PA, Brazil. Emission rates are calculated from the end of a 210 sec period of leaf illumination at 1000 PAR. Max emission per species is provided (tabular), as well as the distributions of emissions from all leaf measurements (graphical). Validation measurements by GC-MS from this study and published estimates for isoprene (I) and monoterpenes (M) from a previous study at the same site are provided for shared species. Genus level data from literature is provided in units of proportion of measured species that emit isoprene (*p*IE) along with the number of species measured (N spp). Numbered references: [1] Present study GC-MS validation measurements; [2] (Bracho-Nunez et al., 2013); [3] (C. Geron et al., 2002); [4] (P. Harley et al., 2004); [5] (K. J. Jardine, Zorzanelli, Gimenez, Oliveira Piva, et al., 2020); [6] (Klinger et al., 2002); [7] (Klinger et al., 1998); [8] (Taylor et al., 2018); [9] (Varshney & Singh, 2003).

This emissions inventory increases tropical species coverage by approximately 10% relative to the literature compilation by Taylor *et al*. (2018). Such inventory datasets contribute to research on the phylogenetic distribution of VI emissions (P. C. Harley et al., 1999; Russell K. Monson et al., 2013), and can be used to refine the ‘emitter fractions’ (proportion of emitting leaf area in a forest) used in emissions and atmospheric chemistry models (C. D. Geron et al., 1994; A. Guenther, 1997). The ability to monitor sensor drift and leaf emission dynamics in real time, and to immediately process emission data without waiting for lab results, allowed the informed adjustment of sampling efforts in the field to ensure adequate coverage of sampling targets and the resolution of any questionable results. We observed detectable (> 0.4 nmol m^-2^ s^-1^), light-dependent emissions from 67% of sampled species. It is likely that emission rates lower than ca. 1 nmol m^-2^ s^-1^ comprise a diversity of compounds, some of which may be passively released by rising temperature when the leaf is illuminated (A. Guenther et al., 1995; Laothawornkitkul et al., 2009). However, measurements subsequent to this campaign have shown that careful measurement of highly aromatic leaves that lack the capacity for light-dependent VI emissions (e.g., *Retrophyllum, Myrcia*, and *Zanthoxylum spp*.) do not produce detectable volatiles during a standard PORCO measurement (data not shown). Emission rates >1 nmol m^-2^ s^-1^ are more likely to be predominantly composed of isoprene, and less commonly, smaller amounts of light-dependent monoterpenes (Harrison et al., 2013; K. J. Jardine, Zorzanelli, Gimenez, Oliveira Piva, et al., 2020; Keller & Lerdau, 1999; Kuhn et al., 2002). All light-dependent VI may play a physiological role similar to isoprene in the leaf, either as antioxidants or as signaling mechanisms for thermal defenses (K. J. Jardine et al., 2017; Vickers, Gershenzon, et al., 2009; Zuo et al., 2019). We observed strong light-dependent emissions (>1 nmol m^-2^ s^-1^) from 59% of sampled species. Consistent with the findings of Taylor et al. (2018), VI emissions show some taxonomic consistency in Table 1. Emission rates ranged from consistently less than 1 nmol m^-2^ s^-1^ in the Lauraceae family, to frequently exceeding 20 nmol m^-2^ s^-1^ in the genus *Protium*. The highest observed emission rate was from an *Eschweilera* species, 28.5 nmol m^-2^ s^-1^.

#### 6.3 Validation of field measurements

Visual analysis of processed leaf-emissions data demonstrate background signal noise to rarely exceed ± 0.2 nmol m^-2^ s^-1^. All emission curves exceeding 0.4 nmol m^-2^ s^-1^ were distinctly different from background fluctuations by 210 sec of illumination (Fig. 13). We therefore assign a conservative uniform detection limit of 0.4 nmol m^-2^ s^-1^ to the inventory dataset (Table 1).

**Figure 13:**
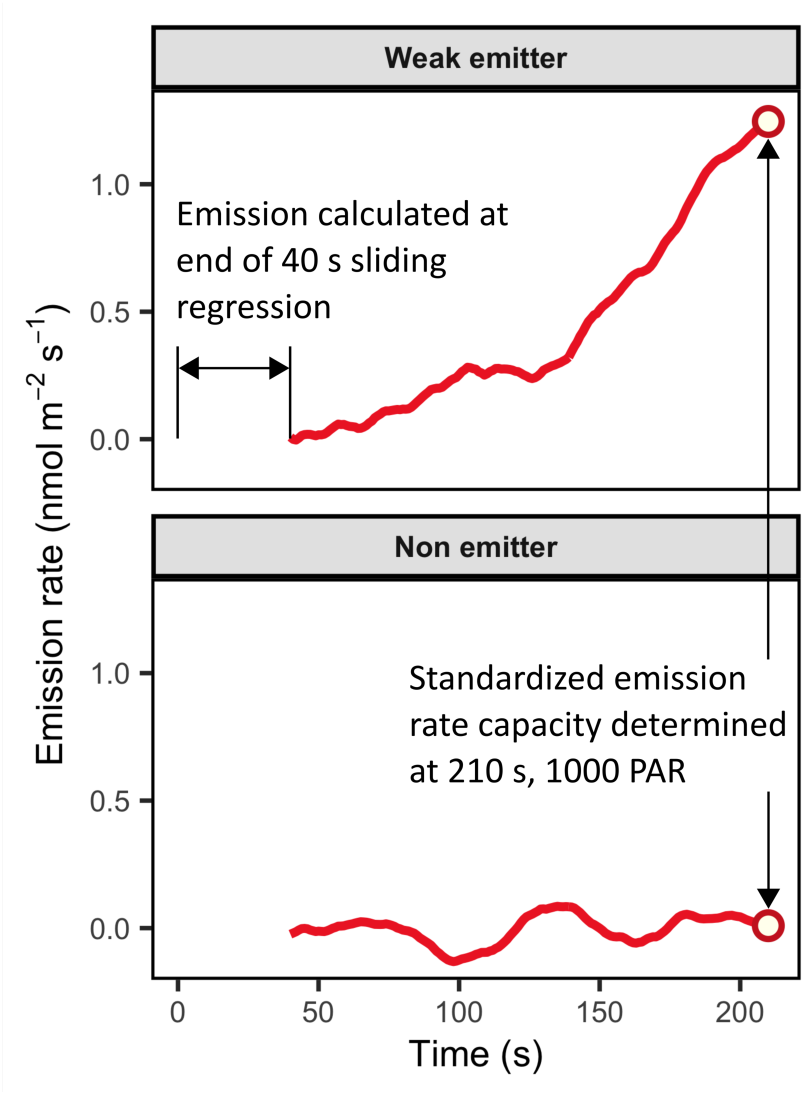
Leaf emission rates are calculated as non-steady-state fluxes based on concentration changes determined by a 40 sec sliding-window regression. Background signal noise produces noise in emission estimates around ± 0.2 nmol m^-2^ s^-1^. Emission rates of 0.4 nmol m^-2^ s^-1^ by 210 sec of measurement time are clearly differentiable from zero.

Much of the variation in emission rates can be expected to occur at the scale of individual branches (Funk et al., 2006; P. Harley et al., 1996; R K Monson & Fall, 1989), which largely determine the light and thermal microenvironment to which leaves are acclimated (Chazdon & Fetcher, 1984; Ü Niinemets et al., 2010). Accordingly, we observed that where within-species variation was highest (Table 1), species data was amalgamated from measurements of multiple distinct trees or branches from different light environments. We tested whether PORCO measurements are sufficiently consistent to detect ecologically driven variation at the branch scale, as distinguished from leaf level variation and detection precision. We found that between-branch variation (standard deviation) in mean emission rates exceeded within-branch (between-leaf) variation by a factor of 1.6 (t-test, *p* << 0.001, Fig. S7), demonstrating that PORCO is able to detect ecologically driven variation in leaf emission rates.

Emission measurements using PORCO show good agreement with other data sources for the same species and genera (Table 1). Independent data was available at the species-level for ten of our measured species. These included our own validation measurements from several of the same individual trees, in which leaves were acclimated to 1000 PAR and 30°C until photosynthetic stability was achieved (approximately 2 min) in a Walz photosynthesis chamber head (Heinz Walz GmbH, Effeltrich, Germany), followed by 10 min of subsampling cuvette gas onto adsorbent cartridges for subsequent analysis by GC-MS, similar to the methods of Jardine et al. (2015). Seven of our measured species were previously measured at the same site by Harley et al. (2004), using a combination of controlled and uncontrolled cuvettes for sampling onto adsorbent cartridges and analysis by gas chromatography. PORCO measurements agreed with independent data sources for all ten shared species in terms of detectable emissions (5 species) versus emissions below detection limit (BDL, 5 species), except *E. uncinatum*, reported as BDL by Harley et al. (2004), but confirmed as an emitter by independent measurements on multiple individuals during this campaign. PORCO measurements also agreed well with published data at the genus-level (Table 1), obtained from a literature compilation of tropical isoprene emission measurements (Taylor et al., 2018, 2019) (Table S2). For species with weak or undetected emissions by PORCO, a low proportion of species from the corresponding genus (“congeners”) were reported as emitters in the literature. Conversely, stronger emissions corresponded to a higher proportion of congeners reported as emitters in the literature (Table 1).

Likely due to the low detection limit achieved with PORCO, we report a higher proportion of species with light-dependent VI emissions (67% above detection limit; 59% > 1 nmol m^-2^ s^-1^) compared the inventory at the same site by Harley et al. (2004) (46% of 26 species, after correcting their reporting of *E. uncinatum* from non-emitter to emitter). Our proportion of detectable emissions closely matches a recent inventory of Amazonian trees based entirely on GC-MS measurements, reporting light-dependent VI emissions from 67% of species (K. J. Jardine, Zorzanelli, Gimenez, Oliveira Piva, et al., 2020). Of the three emitting species independently quantified by GC-MS and PORCO, one emitted only isoprene, one a mixture of isoprene and alpha-pinene (a monoterpene), and one entirely alpha-pinene (Table 1). Estimates of emission rates from the exclusive isoprene emitter were nearly identical (PORCO=18.9, GC-MS=17.8). PORCO measurements increasingly overestimated emissions with an increasing proportion of alpha-pinene from the other two species. Because PIDs have different sensitivities to different compounds (see Section *2*.*1: PID sensitivities, control, and calibration*), we report PORCO measurements calibrated as isoprene-equivalents. The relative responsivity of the ppbRAE 3000 PID to alpha-pinene is reported to be twice that of isoprene (RAE Systems Inc., 2013a), explaining our overestimates of monoterpene emissions when reported as isoprene-equivalents. Although strong, light-dependent monoterpene emissions have been reported from several species (K. J. Jardine et al., 2017; Kuhn et al., 2002), calibration relative to isoprene is the safest bet for a non-distinguishing sensor, as isoprene is demonstrated to be the dominant VI emitted from plants in the tropics (Alves et al., 2016; K. J. Jardine, Zorzanelli, Gimenez, Oliveira Piva, et al., 2020; Sarkar et al., 2020; Yanez-Serrano et al., 2015) and worldwide (Alex Guenther, 2013).

## Results

### Unexpected vertical profile of emission capacities in a tropical forest

If VI emissions are linked to adaptations for leaf thermal tolerance (Sharkey et al., 2008; Taylor et al., 2019; Vickers, Gershenzon, et al., 2009), then species with high emission capacities can be expected to congregate in the upper canopy where light availability and solar heating of leaves is greatest (Ülo Niinemets et al., 2010). Results from our field inventory of tropical trees reveal an unexpected vertical distribution, with the highest emission capacities (Fig. 14a) and the greatest proportion of emitting species (Fig. 14b) found in the mid-canopy region, and surprisingly high emissions found among the smallest trees. We cannot attribute this trend to an over-sampling of canopy species at sub-canopy positions, because we sampled most trees at sizes within the maximum size class for their species. For example, *Protium* species are mid-canopy dominants, never exceeding 45 cm diameter at breast height at our site, and were among the strongest emitters. Two species in the Violaceae family showed consistently high emissions, and they are exclusively understory specialists, never exceeding 10 cm diameter and often found in shaded environments. The strongest observed emission rate was from an 18.8 m tall *Eschweilera sp*. (less than half the height of the tallest trees in this forest) in an open, highly sun exposed crown environment, while similarly exposed canopy-dominant species (*Erisma* and *Tachigali*) showed moderate emissions (Table 1).

**Figure 14:**
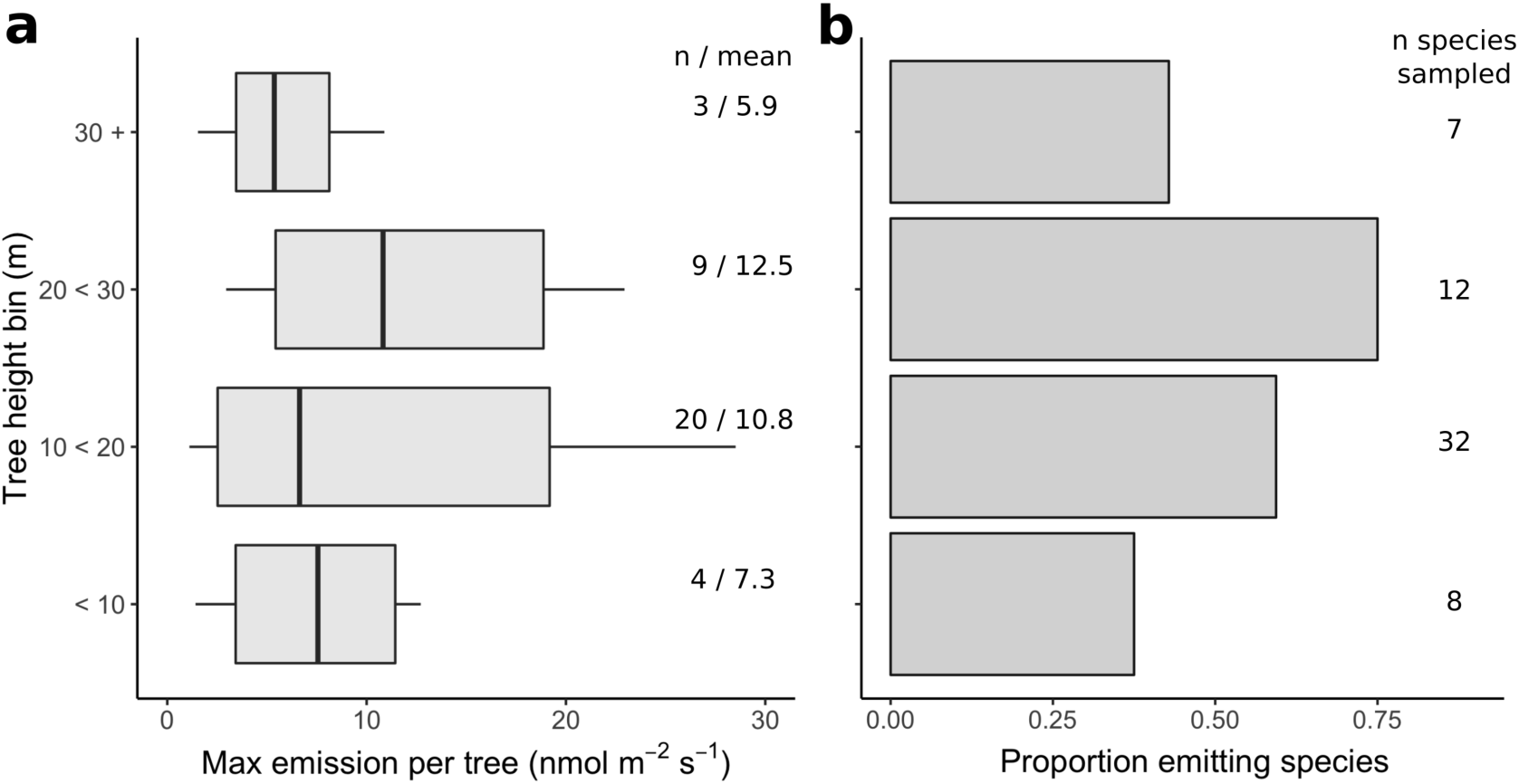
Standardized leaf-level emission rates (3.5 min at 1000 PAR) reach peak values in mid-canopy trees. Boxplots in (a) show the highest emission rate measured per tree (typically 1-3 leaves measured) among trees binned by height. Strong emissions are even observed in the understory. Bars in (b) show the proportion of species with emission rates >1 nmol m^-2^ s^-1^, binned by the maximum tree height among measured individuals of each species.

This vertical distribution of emission capacities suggests a more nuanced view of the adaptive significance of VI emissions for tropical trees. As an on-demand trait, VI emission may be adaptive for temporally dynamic thermal environments (Sharkey et al., 2008), rather than chronically hot ones. In the mid-canopy, leaves experience alternating shade and direct sun exposure as the sun traverses overhead canopy material and gaps. This view presents a challenge for modeling emissions from tropical forests. Forest emissions will depend on the influence of forest structure on dynamic sub-canopy light environments (Ülo Niinemets et al., 2010; Smith et al., 2019; Way & Pearcy, 2012), and cannot be represented as simple steady-state responses to canopy surface conditions. It is worth further exploring how such subtleties in the ecological context of emissions relate to other emerging, nuanced views of the role of VI in dynamic regulation of gene expression (Lantz, Allman, et al., 2019; Zuo et al., 2019) and growth-defense tradeoffs in species’ ecological strategies (Russell K. Monson et al., 2021).

## Discussion

### Contrasting PORCO to commercial instruments: advantages and limitations

PORCO adaptations enable unprecedented in situ detection of leaf volatile emissions with the precision and detection limits required to unambiguously resolve physiological and ecological drivers of emissions from terrestrial vegetation. Compared to other in situ sampling methods, PORCO fills a unique niche with its particular balance of detection capabilities. The ideal detection system would feature: portability; online (real-time) data detection and display; precision detection with low detection limits; leaf environmental control; and the capability to distinguish between gas species. The sections above demonstrate PORCO’s portability and precision online detection capabilities. Its partial leaf environmental control and non-distinguishing sensor represent tradeoffs relative to other methods.

Both precision online detection and compound differentiation can be achieved with a proton transfer reaction mass spectrometer (PTR-MS), but this system is very difficult to mobilize to the field, requiring an improvised laboratory setting (e.g., Alves *et al*., 2016; Sarkar *et al*., 2020). The Fast Isoprene Sensor (A. B. Guenther & Hills, 1998; Hanson & Sharkey, 2001) is more robust and portable than a PTR-MS, and provides online detection of isoprene specifically, but requires AC power and large oxygen tanks to generate ozone for the sensor reaction chamber. PIDs used ‘out of the box’ (not adapted) provide portable online detection, but have poor detection limits due to low signal:noise ratios. Such use requires large, environmentally uncontrollable cuvettes that can enclose many leaves in order to obtain high gas concentrations and overcome sensor noise. The best alternative method currently is to use small adsorbent cartridges to sample gas from the exhaust of commercial leaf cuvettes of photosynthesis instruments that have excellent control of leaf environmental conditions, including light, temperature, CO_2_, and humidity (e.g., Jardine *et al*., 2020a, 2020b). Drawbacks include offline detection (samples are sent to a mass spectrometry lab for analysis), risk of signal degradation during sample transport, expense of cartridges (several hundred USD each), and the limitation of sample sizes to available cartridges and analysis turnaround times.

PORCO leaf cuvettes optimize detection precision and lower-limits, while providing precise light control. However, tight temperature control in larger cuvettes is a challenge. The LED panels warm the air by about 4 °C above ambient by 3.5 minutes. PORCO can alternatively sample from commercial leaf cuvettes, combining precise emission detection with better leaf environmental control (see Section *2*.*2: PORCO-PID validation against Fast Isoprene Sensor*). Drawbacks include an increase in lower detection limit (see Section *3*.*1: Cuvette design*), and the need to obtain, carry, and maintain a second instrument system in the field. The inability to distinguish between compounds with a PID sensor is compensated by the fact that the vast majority of light-dependent emissions from angiosperm tree leaves are isoprene (Alex Guenther, 2013). A non-distinguishing sensor can be considered an advantage in some cases, given that it is capable of detecting a wide range of VI, many of which may play similar physiological roles in the leaf (Vickers, Gershenzon, et al., 2009). The advantages of PORCO can be combined with an ability to distinguish compounds by simultaneously sampling onto adsorbent cartridges from the partially ionized PID exhaust (see Fig. S3), or separately from the leaf cuvettes.

## Conclusion

Despite the well demonstrated critical roles of volatile isoprenoid emissions in plant physiology and atmospheric chemistry, the field research required to bridge the scales from leaf to atmosphere has been held back by methodological limitations. The adaptations presented in PORCO resolve key limitations, allowing an increased rate of species sampling and the in situ measurement precision required to access exciting new questions in ecology, evolution, and atmospheric science. For example, modeling forest emissions requires that we understand the landscape distributions of emitting species (A. Guenther, 1997; Sharkey & Monson, 2014). The present and future distributions of heat-tolerant emitters may also be a key determinant of the sensitivity of forest carbon uptake to climate warming (Feeley et al., 2020; Smith et al., 2020). Limited data suggests that the enhanced thermal tolerance of isoprene emitting species shapes their distributions across tropical landscapes and through time (Taylor et al., 2019), but the quantified uncertainty in emitter distributions is high due to limited species sampling (Taylor et al., 2018). A conspicuous loss of emitters toward dry climates (Klinger et al., 1998; Taylor et al., 2018) remains enigmatic, and may be linked to unexplored coordination between emissions and leaf carbon-economic and hydraulic strategies (Taylor et al., 2018).

Field campaigns will enable targeted hypothesis testing, and species inventories such as reported here (Table 1) increase the potential to correlate emissions with other published datasets describing species thermal tolerance strategies and other ecophysiological traits (Perez et al., 2018; Perez & Feeley, 2020; I. J. Wright et al., 2004; S. J. Wright et al., 2010) or demographic responses to climate (Duque et al., 2015; Enquist & Enquist, 2011; Fadrique et al., 2018). While the diversity of tropical forests is overwhelming (Slik et al., 2015), targeted sampling can resolve the phylogenetic (evolutionary) distribution of emissions among species (P. C. Harley et al., 1999; Russell K. Monson et al., 2013), which will allow for more informed inference of unmeasured species (Taylor et al., 2018). The vertical profile of emission capacities presented here suggests that canopy structure may mediate forest emissions by determining spatiotemporal variation in light environments of the middle and low canopy (e.g., Smith *et al*., 2019). The generality of this vertical distribution and the mechanisms driving it should be explored further. PORCO has the portability and speed of online detection required to observe how emissions respond to temporally dynamic sun exposure beneath the canopy surface. The ease of deployment of PORCO can enable repeat measurements to disentangle the effects of leaf age and environment (Alves et al., 2014; Ü Niinemets et al., 2010) that contribute to a ‘cryptic phenology’ of emissions from evergreen forests (Albert et al., 2019; Alves et al., 2016, 2018). Understanding emission responses to leaf age, macro and microenvironmental variation, and the mechanisms underpinning species distributions will aid in scaling efforts based on physiological emissions models (Morfopoulos et al., 2014; Unger et al., 2013) and remote sensing (Zheng et al., 2015, 2017) in order to better resolve the role of emissions in biosphere-anthroposphere-climate interactions (Unger et al., 2017).

PORCO adaptations could also benefit applied research. VI-emitting crops may be more resistant to drought and ozone, but at a cost to growth rate (Ryan et al., 2014; Vickers, Possell, et al., 2009). With the efficient field sampling enabled by PORCO, crop strains could be monitored for selection of optimum emission rates. The emission rates of urban trees can be monitored to understand their impacts on air quality for human health (Montoya et al., 2020; Purser, Heal, et al., 2020; Wang et al., 2013; Zheng et al., 2020). Our new system brings the laboratory to the leaf, opening a new frontier in research toward a mechanistic understanding of volatile isoprenoid emissions from plants.

## Supporting information

Taylor-2021-Supplemental

## Acknowledgments

Financial support for this study was provided to: T.C.T., W.W., and S.R.S. by grant NSF-PIRE #OISE-0730305 and the DOE GoAmazon project award #3002937712; T.C.T. by NSF grant #NSF-PRFB-1711997. We thank Dr. Kolby Jardine for advising in the early development stages. We thank Prof. Russel K. Monson for advising and access to validation instrumentation. We thank the botanical personnel at Herbario-IAN, Embrapa, in Belém, PA, Brazil, for identifying our plant specimens free of charge to support T.C.T.’s dissertation research. T.C.T. thanks Tiago Martins for his dedicated, reliable help in the field, as well as Mick Eltringham and Neill Prohaska for canopy access assistance. We thank Tucson Store Fixtures for careful construction of the leaf cuvettes according to our designs. We thank the University of Arizona’s Biosphere 2 facility and Professor Julio Tota at the Universidade Federal do Oeste do Pará, Brazil for laboratory space during instrument development. Parts of this article content were published online previously in the Ph.D. dissertation of Tyeen C. Taylor (Taylor, 2017). The submitted form of this article was posted as a pre-print on bioRxiv.

## References

Albert, L. P., Smith, M. N., Bryant, C. C., Prohaska, N., Taylor, T. C., Martins, G. A., et al. (2019). Cryptic phenology in plants?: Case studies, implications, and recommendations, (June), 3591–3608. https://doi.org/10.1111/gcb.14759

Alves, E. G., Harley, P., Gonçalves, J. F. D. C., Eduardo, C., & Jardine, K. (2014). Effects of light and temperature on isoprene emission at different leaf developmental stages of Eschweilera coriacea in central Amazon. Acta Amazonica, 44(1), 9–18. https://doi.org/10.1590/S0044-59672014000100002

Alves, E. G., Jardine, K., Tota, J., Jardine, A., Maria Yãnez-Serrano, A., Karl, T., et al. (2016). Seasonality of isoprenoid emissions from a primary rainforest in central Amazonia. Atmospheric Chemistry and Physics, 16(6), 3903–3925. https://doi.org/10.5194/acp-16-3903-2016

Alves, E. G., Tóta, J., Turnipseed, A., Guenther, A. B., Bustillos, J. O. W. V., Santana, R. A., et al. (2018). Leaf phenology as one important driver of seasonal changes in isoprene emission in central Amazonia. Biogeosciences Discussions, (March), 4–5. https://doi.org/10.5194/bg-15-4019-2018

Behnke, K., Ehlting, B., Teuber, M., Bauerfeind, M., Louis, S., Hänsch, R., et al. (2007). Transgenic, non-isoprene emitting poplars don’t like it hot. The Plant Journal, 51(3), 485–499. https://doi.org/10.1111/j.1365-313X.2007.03157.x

Boyle, B., Hopkins, N., Lu, Z., Raygoza Garay, J. A., Mozzherin, D., Rees, T., et al. (2013). The taxonomic name resolution service: an online tool for automated standardization of plant names. BMC Bioinformatics, 14, 1–14. https://doi.org/10.1186/1471-2105-14-16

Bracho-Nunez, A., Knothe, N. M., Welter, S., Staudt, M., Costa, W. R., Liberato, M. A. R., et al. (2013). Leaf level emissions of volatile organic compounds (VOC) from some Amazonian and Mediterranean plants. Biogeosciences, 10, 5855–5873. https://doi.org/10.5194/bg-10-5855-2013

Carslaw, K. S., Boucher, O., Spracklen, D. V., Mann, G. W., Rae, J. G. L., Woodward, S., & Kulmala, M. (2010). A review of natural aerosol interactions and feedbacks within the Earth system. Atmospheric Chemistry and Physics, 10(4), 1701–1737. https://doi.org/10.5194/acp-10-1701-2010

Carslaw, K. S., Lee, L. A., Reddington, C. L., Pringle, K. J., Rap, A., Forster, P. M., et al. (2013). Large contribution of natural aerosols to uncertainty in indirect forcing. Nature, 503(7474), 67–71. https://doi.org/10.1038/nature12674

Chazdon, R. L., & Fetcher, N. (1984). Photosynthetic Light Environments in a Lowland Tropical Rain Forest in Costa Rica. Journal of Ecology, 72(2), 553–564. https://doi.org/10.2307/2260066

Duque, A., Stevenson, P. R., & Feeley, K. J. (2015). Thermophilization of adult and juvenile tree communities in the northern tropical Andes. Proceedings of the National Academy of Sciences, 112(34), 10744–10749. https://doi.org/10.1073/pnas.1506570112

Enquist, B. J., & Enquist, C. A. F. (2011). Long-term change within a Neotropical forest: Assessing differential functional and floristic responses to disturbance and drought. Global Change Biology, 17(3), 1408–1424. https://doi.org/10.1111/j.1365-2486.2010.02326.x

Fadrique, B., Báez, S., Duque, Á., Malizia, A., Blundo, C., Carilla, J., et al. (2018). Widespread but heterogeneous responses of Andean forests to climate change. Nature, 564, 207–212. https://doi.org/10.1038/s41586-018-0715-9

Feeley, K. J., Bravo-Avila, C., Fadrique, B., Perez, T. M., & Zuleta, D. (2020). Climate-driven changes in the composition of New World plant communities. Nature Climate Change, 10(10), 965–970. https://doi.org/10.1038/s41558-020-0873-2

Feng, Z., Yuan, X., Fares, S., Loreto, F., Li, P., Hoshika, Y., & Paoletti, E. (2019). Isoprene is more affected by climate drivers than monoterpenes: A meta-analytic review on plant isoprenoid emissions. Plant Cell and Environment, 42(6), 1939– 1949. https://doi.org/10.1111/pce.13535

Fineschi, S., Loreto, F., Staudt, M., & Peñuelas, J. (2013). Diversification of volatile isoprenoid emissions from trees: evolutionary and ecological perspectives. In Ülo Niinemets & R. K. Monson (Eds.), Biology, Controls and Models of Tree Volatile Organic Compound Emissions (pp. 1–20). Springer. https://doi.org/10.1007/978-94-007-6606-8

Fini, A., Brunetti, C., Loreto, F., Centritto, M., Ferrini, F., & Tattini, M. (2017). Isoprene responses and functions in plants challenged by environmental pressures associated to climate change. Frontiers in Plant Science, 8, 1281. https://doi.org/10.3389/FPLS.2017.01281

Funk, J. L., Giardina, C. P., Knohl, A., & Lerdau, M. T. (2006). Influence of nutrient availability, stand age, and canopy structure on isoprene flux in a Eucalyptus saligna experimental forest. Journal of Geophysical Research: Biogeosciences, 111(2), 1– 10. https://doi.org/10.1029/2005JG000085

Geron, C., Guenther, A., Greenberg, J., Loescher, H. W., Clark, D., & Baker, B. (2002). Biogenic volatile organic compound emissions from a lowland tropical wet forest in Costa Rica. Atmospheric Environment, 36, 3793–3802. https://doi.org/10.1016/S1352-2310(02)00301-1

Geron, C., Daly, R., Harley, P., Rasmussen, R., Seco, R., Guenther, A., et al. (2016). Large drought-induced variations in oak leaf volatile organic compound emissions during PINOT NOIR 2012. Chemosphere, 146, 8–21. https://doi.org/10.1016/j.chemosphere.2015.11.086

Geron, C. D., Guenther, A. B., & Pierce, T. E. (1994). An improved model for estimating emissions of volatile organic compounds from forests in the eastern United States. Journal of Geophysical Research, 99(D6), 12773–12791. https://doi.org/10.1029/94JD00246

Guenther, A. (1997). Seasonal and spatial variations in natural volatile organic compound emissions. Ecological Applications, 7(1), 34–45. https://doi.org/10.1890/1051-0761(1997)007[0034:SASVIN]2.0.CO;2

Guenther, A., Hewitt, C. N., Erickson, D., Fall, R., Geron, C., Graedel, T., et al. (1995). A global model of natural volatile organic compound emissions. Journal of Geophysical Research, 100(D5), 8873. https://doi.org/10.1029/94JD02950

Guenther, A. B., & Hills, A. J. (1998). Eddy covariance measurement of isoprene fluxes. Journal of Geophysical Research Atmospheres, 103(D11), 13145–13152. https://doi.org/10.1029/97JD03283

Guenther, Alex. (2013). Biological and Chemical Diversity of Biogenic Volatile Organic Emissions into the Atmosphere. ISRN Atmospheric Sciences, 2013(January), 1–27. https://doi.org/10.1155/2013/786290

Hanson, D. T., & Sharkey, T. D. (2001). Effect of growth conditions on isoprene emission and other thermotolerance-enhancing compounds. Plant, Cell and Environment, 24(9), 929–936. https://doi.org/10.1046/j.1365-3040.2001.00744.x

Hantson, S., Knorr, W., Schurgers, G., Pugh, T. A. M., & Arneth, A. (2017). Global isoprene and monoterpene emissions under changing climate, vegetation, CO2 and land use. Atmospheric Environment, 155, 35–45. https://doi.org/10.1016/j.atmosenv.2017.02.010

Harley, P., Guenther, A., & Zimmerman, P. (1996). Effects of light, temperature and canopy position on net photosynthesis and isoprene emission from sweetgum (Liquidambar styraciflua) leaves. Tree Physiology, 16, 25–32. https://doi.org/10.1093/treephys/16.1-2.25

Harley, P., Vasconcellos, P., Vierling, L., Pinheiro, C. C. D. S., Greenberg, J., Guenther, A., et al. (2004). Variation in potential for isoprene emissions among Neotropical forest sites. Global Change Biology, 10(5), 630–650. https://doi.org/10.1111/j.1529-8817.2003.00760.x

Harley, P. C., Monson, R. K., & Lerdau, M. T. (1999). Ecological and evolutionary aspects of isoprene emission from plants. Oecologia, 118(2), 109–123. https://doi.org/10.1007/s004420050709

Harrison, S. P., Morfopoulos, C., Dani, K. G. S., Prentice, I. C., Arneth, A., Atwell, B. J., et al. (2013). Volatile isoprenoid emissions from plastid to planet. New Phytologist, 197(1), 49–57. https://doi.org/10.1111/nph.12021

Heald, C. L., Henze, D. K., Horowitz, L. W., Feddema, J., Lamarque, J.-F., Guenther, A., et al. (2008). Predicted change in global secondary organic aerosol concentrations in response to future climate, emissions, and land use change. Journal of Geophysical Research, 113(D5), D05211. https://doi.org/10.1029/2007JD009092

Hunt, S. (2003). Measurements of photosynthesis and respiration in plants. Physiologia Plantarum, 117(3), 314–325. https://doi.org/10.1034/j.1399-3054.2003.00055.x

Jardine, A. B., Jardine, K. J., Fuentes, J. D., Martin, S. T., Martins, G., Durgante, F., et al. (2015). Highly reactive light-dependent monoterpenes in the Amazon. Geophysical Research Letters, 42(5), 1576–1583. https://doi.org/10.1002/2014GL062573

Jardine, K. J., Jardine, A. B., Holm, J. A., Lombardozzi, D. L., Negron-juarez, R. I., & Martin, S. T. (2017). Monoterpene ‘thermometer’ of tropical forest-atmosphere response to climate warming. Plant, Cell and Environment, 441–452. https://doi.org/10.1111/pce.12879

Jardine, K. J., Zorzanelli, R. F., Gimenez, B. O., Robles, E., & de Oliveira Piva, L.R. (2020). Development of a portable leaf photosynthesis and volatile organic compounds emission system. MethodsX, 7. https://doi.org/10.1016/j.mex.2020.100880

Jardine, K. J., Zorzanelli, R. F., Gimenez, B. O., Oliveira Piva, L. R. de, Teixeira, A., Fontes, C. G., et al. (2020). Leaf isoprene and monoterpene emission distribution across hyperdominant tree genera in the Amazon basin. Phytochemistry, 175(April), 112366. https://doi.org/10.1016/j.phytochem.2020.112366

Keller, M., & Lerdau, M. (1999). Isoprene emission from tropical forest canopy leaves. Global Biogeochemical Cycles, 13(1), 19–29. https://doi.org/10.1029/1998GB900007

Kim, S., Chen, J., Cheng, T., Gindulyte, A., He, J., He, S., et al. (2020). PubChem in 2021: new data content and improved web interfaces. Nucleic Acids Research, 49(November), 1388–1395. https://doi.org/10.1093/nar/gkaa971

Klinger, L. F., Greenberg, J., Guenther, A., Zimmerman, P., M’Bangui, M., & Kenfack, D. (1998). Patterns in volatile organic compound emissions along a savanna-rainforest gradient in central Africa. Journal of Geophysical Research, 103(D1), 1443–1454. https://doi.org/10.1029/97JD02928

Klinger, L. F., Li, Q. J., Guenther, A. B., Greenberg, J. P., Baker, B., & Bai, J. H. (2002). Assessment of volatile organic compound emissions from ecosystems of China. Journal of Geophysical Research Atmospheres, 107(21). https://doi.org/10.1029/2001JD001076

Kuhn, U., Rottenberger, S., Biesenthal, T., Wolf, A., Schebeske, G., Ciccioli, P., et al. (2002). Isoprene and monoterpene emissions of Amazônian tree species during the wet season: Direct and indirect investigations on controlling environmental functions. Journal of Geophysical Research, 107(D20), 1–13. https://doi.org/10.1029/2001JD000978

Lantz, A. T., Allman, J., Weraduwage, S. M., & Sharkey, T. D. (2019). Isoprene: New insights into the control of emission and mediation of stress tolerance by gene expression. Plant Cell and Environment, 42(10), 2808–2826. https://doi.org/10.1111/pce.13629

Lantz, A. T., Solomon, C., Gog, L., McClain, A. M., Weraduwage, S. M., Cruz, J. A., & Sharkey, T. D. (2019). Isoprene suppression by CO2 is not due to triose phosphate utilization (TPU) limitation. Frontiers in Forests and Global Change, 2(April), 1– 13. https://doi.org/10.3389/ffgc.2019.00008

Laothawornkitkul, J., Taylor, J. E., Paul, N. D., & Hewitt, C. N. (2009). Biogenic volatile organic compounds in the Earth system: Tansley review. New Phytologist, 183(1), 27–51. https://doi.org/10.1111/j.1469-8137.2009.02859.x

Li, Z., & Sharkey, T. D. (2013). Metabolic profiling of the methylerythritol phosphate pathway reveals the source of post-illumination isoprene burst from leaves. Plant, Cell and Environment, 36(2), 429–437. https://doi.org/10.1111/j.1365-3040.2012.02584.x

Martins, M. A. R., Silva, L. P., Ferreira, O., Schröder, B., Coutinho, J. A. P., & Pinho, S. P. (2017). Terpenes solubility in water and their environmental distribution. Journal of Molecular Liquids, 241, 996–1002. https://doi.org/10.1016/j.molliq.2017.06.099

Monson, R K, & Fall, R. (1989). Isoprene emission from aspen leaves: influence of environment and relation to photosynthesis and photorespiration. Plant Physiology, 90(1), 267–74. https://doi.org/10.1104/pp.90.1.267

Monson, R K, Lerdau, M. T., Sharkey, T. D., Schimel, D. S., & Fall, R. (1995). Biological Aspects of Constructing Volatile Organic-Compound Emission Inventories. Atmospheric Environment, 29(21), 2989–3002. https://doi.org/10.1016/1352-2310(94)00360-W

Monson, Russell K., Jones, R. T., Rosenstiel, T. N., & Schnitzler, J. P. (2013). Why only some plants emit isoprene. Plant, Cell and Environment, 36(3), 503–516. https://doi.org/10.1111/pce.12015

Monson, Russell K., Neice, A. A., Trahan, N. A., Shiach, I., McCorkel, J. T., & Moore D.J. P. (2016). Interactions between temperature and intercellular CO2 concentration in controlling leaf isoprene emission rates. Plant Cell and Environment, 39(11), 2404–2413. https://doi.org/10.1111/pce.12787

Monson, Russell K., Weraduwage, S. M., Rosenkranz, M., Schnitzler, J. P., & Sharkey, T. D. (2021). Leaf isoprene emission as a trait that mediates the growth-defense tradeoff in the face of climate stress. Oecologia, (3). https://doi.org/10.1007/s00442-020-04813-7

Montoya, O. L. Q., Niño-Ruiz, E. D., & Pinel, N. (2020). On the mathematical modelling and data assimilation for air pollution assessment in the Tropical Andes. Environmental Science and Pollution Research, 27(29), 35993–36012. https://doi.org/10.1007/s11356-020-08268-4

Morfopoulos, C., Sperlich, D., Peñuelas, J., Filella, I., Llusià, J., Medlyn, B. E., et al. (2014). A model of plant isoprene emission based on available reducing power captures responses to atmospheric CO2. The New Phytologist, 203(1), 125–39. https://doi.org/10.1111/nph.12770

Niinemets, Ü, Arneth, A., Kuhn, U., Monson, R. K., Peñuelas, J., & Staudt, M. (2010). The emission factor of volatile isoprenoids: Stress, acclimation, and developmental responses. Biogeosciences, 7(7), 2203–2223. https://doi.org/10.5194/bg-7-2203-2010

Niinemets, Ülo, Copolovici, L., & Hüve, K. (2010). High within-canopy variation in isoprene emission potentials in temperate trees: Implications for predicting canopy-scale isoprene fluxes. Journal of Geophysical Research: Biogeosciences, 115(4), 1– 19. https://doi.org/10.1029/2010JG001436

Padhy, P. K., & Varshney, C. K. (2005). Isoprene emission from tropical tree species. Environmental Pollution, 135, 101–109. https://doi.org/10.1016/j.envpol.2004.10.003

Perez, T. M., & Feeley, K. J. (2020). Photosynthetic heat tolerances and extreme leaf temperatures. Functional Ecology, 34(11), 2236–2245. https://doi.org/10.1111/1365-2435.13658

Perez, T. M., Valverde-barrantes, O., Bravo, C., Hogan, J. A., Pardo, C. J., Taylor, T. C., et al. (2018). Botanic gardens are an untapped resource for studying the functional ecology of tropical plants. Philosophical Transactions of the Royal Society of London. Series B, Biological Sciences, 374(1763), 20170390. https://doi.org/10.1098/rstb.2017.0390

Purser, G., Heal, M. R., White, S., Morison, J. I. L., & Drewer, J. (2020). Differences in isoprene and monoterpene emissions from cold-tolerant eucalypt species grown in the UK. Atmospheric Pollution Research, 11(11), 2011–2021. https://doi.org/10.1016/j.apr.2020.07.022

Purser, G., Drewer, J., Heal, M., Sircus, R., Dunn, L., & Morison, J. (2020). Isoprene and monoterpene emissions from alder, aspen and spruce short rotation forest plantations in the UK. Biogeosciences Discussions, (December), 1–52. https://doi.org/10.5194/bg-2020-437

R Core Team. (2020). R: A language and environment for statistical computing. Vienna, Austria: R Foundation for Statistical Computing. Retrieved from https://www.r-project.org/.

RAE Systems Inc. (2013a). Technical Note TN-106 Correction Factors, Ionization Energies, and Calibration Characteristics.

RAE Systems Inc. (2013b). Technical Note TN-165: Combating Drift in Portable and Fixed PIDs.

RAE Systems Inc. (2013c). The PID Handbook. Charlotte, NC: Honeywell International Inc.

Rasmussen, R., & Went, F. W. (1964). Volatile organic matter of plant origin in the atmosphere. Science, 144(3618), 566. https://doi.org/10.1126/science.144.3618.566-a

Rice, A. H., Pyle, E. H., Saleska, S. R., Hutyra, L., Keller, M., Camargo, P. B. De, et al. (2004). Carbon Balance and Vegetation Dynamics in an Old-Growth Amazonian Forest. Ecological Applications, 14(4), S55–S71. https://doi.org/10.1890/02-6006

Rieksta, J., Li, T., Junker, R. R., Jepsen, J. U., Ryde, I., & Rinnan, R. (2020). Insect Herbivory Strongly Modifies Mountain Birch Volatile Emissions. Frontiers in Plant Science, 11(October). https://doi.org/10.3389/fpls.2020.558979

Rinnan, R., Iversen, L. L., Tang, J., Vedel-Petersen, I., Schollert, M., & Schurgers, G. (2020). Separating direct and indirect effects of rising temperatures on biogenic volatile emissions in the Arctic. Proceedings of the National Academy of Sciences of the United States of America, 117(51), 32476–32483. https://doi.org/10.1073/pnas.2008901117

Ryan, A. C., Hewitt, C. N., Possell, M., Vickers, C. E., Purnell, A., Mullineaux, P. M., et al. (2014). Isoprene emission protects photosynthesis but reduces plant productivity during drought in transgenic tobacco (Nicotiana tabacum) plants. New Phytologist, 201(1), 205–216. https://doi.org/10.1111/nph.12477

Sanadze, G. A. (2004). Biogenic isoprene (a review). Russian Journal of Plant Physiology, 51(6), 729–741. https://doi.org/10.1023/B:RUPP.0000047821.63354.a4

Sarkar, C., Guenther, A. B., Park, J., Seco, R., Alves, E., Batalha, S., et al. (2020). PTR-TOF-MS eddy covariance measurements of isoprene and monoterpene fluxes from an eastern Amazonian rainforest. Atmospheric Chemistry and Physics, 20, 7179– 7191. https://doi.org/10.5194/acp-20-7179-2020

Seco, R., Holst, T., Sillesen Matzen, M., Westergaard-Nielsen, A., Li, T., Simin, T., et al. (2020). Volatile organic compound fluxes in a subarctic peatland and lake. Atmospheric Chemistry and Physics, 20(21), 13399–13416. https://doi.org/10.5194/acp-20-13399-2020

Sharkey, T. D., & Monson, R. K. (2014). The future of isoprene emission from leaves, canopies and landscapes. Plant, Cell and Environment, 37(8), 1727–1740. https://doi.org/10.1111/pce.12289

Sharkey, T. D., & Monson, R. K. (2017). Isoprene research - 60 years later, the biology is still enigmatic. Plant Cell and Environment, 40(9), 1671–1678. https://doi.org/10.1111/pce.12930

Sharkey, T. D., & Yeh, S. (2001). Isoprene emission from plants. Annual Review of Plant Physiology and Plant Molecular Biology, 52, 407–436. https://doi.org/10.1093/aob/mcm240

Sharkey, T. D., Wiberley, A. E., & Donohue, A. R. (2008). Isoprene emission from plants: Why and how. Annals of Botany, 101(1), 5–18. https://doi.org/10.1093/aob/mcm240

Singsaas, E. L., Lerdau, M., Winter, K., & Sharkey, T. D. (1997). Isoprene Increases Thermotolerance of Isoprene-Emitting Species. Plant Physiology, 115(4), 1413– 1420. https://doi.org/10.1104/pp.115.4.1413

Slik, J. W. F., Arroyo-Rodriguez, V., Shin-Ichiro, A., Alvarez-loayza, P., Alves, L. F., Ashton, P., et al. (2015). An estimate of the number of tropical tree species. Proceedings of the National Academy of Sciences, 112(34), 7472–7477. https://doi.org/10.1073/pnas.1512611112

Smith, M. N., Stark, S. C., Taylor, T. C., Ferreira, M. L., de Oliveira, E., Restrepo-Coupe, N., et al. (2019). Seasonal and drought related changes in leaf area profiles depend on height and light environment in an Amazon forest. New Phytologist, 222(3), 1284–1297. https://doi.org/10.1111/nph.15726

Smith, M. N., Taylor, T. C., van Haren, J., Rosolem, R., Restrepo-Coupe, N., Adams, J., et al. (2020). Empirical evidence for resilience of tropical forest photosynthesis in a warmer world. Nature Plants, 6(10), 1225–1230. https://doi.org/10.1038/s41477-020-00780-2

Tani, A., & Mochizuki, T. (2021). Review: Exchanges of volatile organic compounds between terrestrial ecosystems and the atmosphere. Journal of Agricultural Meteorology, 77(1), 66–80. https://doi.org/10.2480/agrmet.D-20-00025

Taylor, T. C. (2017). Ph.D. Thesis: Leaf Volatile Emissions Structure Tree Community Assembly and Mediate Climate Feedbacks in Tropical Forests. University of Arizona. University of Arizona. Retrieved from http://hdl.handle.net/10150/623061

Taylor, T. C., Mcmahon, S. M., Smith, M. N., Boyle, B., Violle, C., Haren J. Van, et al. (2018). Isoprene emission structures tropical tree biogeography and community assembly responses to climate. New Phytologist, 220, 435–446. https://doi.org/10.1111/nph.15304

Taylor, T. C., Smith, M. N., Slot, M., & Feeley, K. J. (2019). The capacity to emit isoprene differentiates the photosynthetic temperature responses of tropical plant species. Plant, Cell & Environment, 42(8), 2448–2457. https://doi.org/10.1111/pce.13564

Unger, N. (2014). On the role of plant volatiles in anthropogenic global climate change. Geophysical Research Letters, 41(23), 8563–8569. https://doi.org/10.1002/2014GL061616

Unger, N., Harper, K., Zheng, Y., Kiang, N. Y., Aleinov, I., Arneth, A., et al. (2013). Photosynthesis-dependent isoprene emission from leaf to planet in a global carbon-chemistry-climate model. Atmospheric Chemistry and Physics, 13(20), 10243– 10269. https://doi.org/10.5194/acp-13-10243-2013

Unger, N., Yue, X., & Harper, K. L. (2017). Aerosol climate change effects on land ecosystem services. Faraday Discussions, 200, 121–142. https://doi.org/10.1039/c7fd00033b

Varshney, C. K., & Singh, A. P. (2003). Isoprene emission from Indian trees. Journal of Geophysical Research, 108(D24), ACH24-1–7. https://doi.org/10.1029/2003JD003866

Vickers, C. E., Gershenzon, J., Lerdau, M. T., & Loreto, F. (2009). A unified mechanism of action for volatile isoprenoids in plant abiotic stress. Nature Chemical Biology, 5(5), 283–291. https://doi.org/10.1038/nchembio.158

Vickers, C. E., Possell, M., Cojocariu, C. I., Velikova, V. B., Laothawornkitkul, J., Ryan, A., et al. (2009). Isoprene synthesis protects transgenic tobacco plants from oxidative stress. Plant, Cell and Environment, 32(5), 520–531. https://doi.org/10.1111/j.1365-3040.2009.01946.x

Wang, J. L., Chew, C., Chang, C. Y., Liao, W. C., Lung, S. C. C., Chen, W. N., et al. (2013). Biogenic isoprene in subtropical urban settings and implications for air quality. Atmospheric Environment, 79, 369–379. https://doi.org/10.1016/j.atmosenv.2013.06.055

Way, D. A., & Pearcy, R. W. (2012). Sunflecks in trees and forests: From photosynthetic physiology to global change biology. Tree Physiology, 32(9), 1066–1081. https://doi.org/10.1093/treephys/tps064

Wickham, H. (2016). ggplot2: Elegant Graphics for Data Analysis. New York: Springer-Verlag.

Wright, I. J., Reich, P. B., Westoby, M., Ackerly, D. D., Baruch, Z., Bongers, F., et al. (2004). The worldwide leaf economics spectrum. Nature, 428(6985), 821–827. https://doi.org/10.1038/nature02403

Wright, S. J., Kitajima, K., Kraft, N. J. B., Reich, P. B., Wright, I. J., Bunker, D. E., et al. (2010). Functional traits and the growth-mortality trade-off in tropical trees. Ecology, 91(12), 3664–74. https://doi.org/10.1890/09-2335.1

Yanez-Serrano, A. M., Nölscher, a C., Williams, J., Wolff, S., Alves, E., Martins, G. a, & Bourtsoukidis, E. (2015). Diel and seasonal changes of biogenic volatile organic compounds within and above an Amazonian rainforest, (x), 3359–3378. https://doi.org/10.5194/acp-15-3359-2015

Yáñez-serrano, A. M., Bourtsoukidis, E., Alves, E. G., Bauwens, M., Stavrakou, T., Llusià, J., et al. (2020). Amazonian biogenic volatile organic compounds under global change. Global Change Biology, (May), 1–30. https://doi.org/10.1111/gcb.15185

Zheng, Y., Unger, N., Barkley, M. P., & Yue, X. (2015). Relationships between photosynthesis and formaldehyde as a probe of isoprene emission. Atmospheric Chemistry and Physics, 15(15), 8559–8576. https://doi.org/10.5194/acp-15-8559-2015

Zheng, Y., Unger, N., Tadić, J. M., Seco, R., Guenther, A. B., Barkley, M. P., et al. (2017). Drought impacts on photosynthesis, isoprene emission and atmospheric formaldehyde in a mid-latitude forest. Atmospheric Environment, 167, 190–201. https://doi.org/10.1016/j.atmosenv.2017.08.017

Zheng, Y., Thornton, J. A., Lee Ng, N., Cao, H., Henze, D. K., McDuffie, E. E., et al. (2020). Long-term observational constraints of organic aerosol dependence on inorganic species in the southeast US. Atmospheric Chemistry and Physics, 20(21), 13091–13107. https://doi.org/10.5194/acp-20-13091-2020

Zuo, Z., Weraduwage, S. M., Lantz, A. T., Sanchez, L. M., Weise, S. E., Wang, J., et al. (2019). Isoprene acts as a signaling molecule in gene networks important for stress responses and plant growth. Plant Physiology, 180(1), 124–152. https://doi.org/10.1104/pp.18.01391

