## Supplementary material for "A new field instrument for leaf volatiles reveals an unexpected vertical profile of isoprenoid emission capacities in a tropical forest": Taylor-2021-Supplemental

**List of authors**

Tyeen C. Taylor[1,2], Wit T. Wisniewski[1], Eliane G. Alves[3], Raimundo C. de Oliveira[4], Scott R. Saleska[1]

**Affiliations**

[1]Department of Ecology and Evolutionary Biology, University of Arizona, Tucson, AZ, USA.

[2]Biology Department, University of Miami, Miami, FL, USA

[3]Department of Biogeochemical Processes, Max Planck Institute for Biogeochemistry, Jena, Germany.

[4]Embrapa Amazônia Oriental, Santarém, PA, Brazil

**Correspondence**

Dr. Tyeen C. Taylor:

**Table S1:** Major components of the PORCO system.

| Component | Source | Purpose |
|---|---|---|
| Detector | Honeywell International Inc., Charlotte, NC, USA | ppbRAE-3000 photoionization detector, the gas detection instrument used in PORCO. |
| Mass flow controllers | Alicat Scientific, Tucson, AZ, USA | A 0-500 and a 0-1000 sccm mass flow controller for mixing calibration gas and zero-air, and controlling zero-air flow rates to leaf cuvettes |
| Refillable hydrocarbon trap | Restek Corporation, Bellefonte, PA, USA | Purifying ambient air of hydrocarbons, i.e. produces 'zero air'. |
| Air pump | Parker Hannifin, Hollis, NH, USA | CTS Micro diaphragm pump. Air pump for positive pressure flow of zero-air for calibration and sampling. |
| iButtons | Maxim Integrated, San Jose, CA, USA | Datalogging temperature and humidity |
| Gas fittings and tubing | Swagelok Southwest Co., Phoenix, AZ, USA | Stainless steel fittings and valves, and PFA tubing for primary flow pathways. |

| | | |
|---|---|---|
| LED-Red | Cree Inc, Durham, NC, USA | Cree Xlamp "Photo Red" surface mount high power LED for providing photosynthetically active radiation to leaf. |
| LED-Blue | Cree Inc, Durham, NC, USA | Cree Xlamp "Royal Blue" surface mount high power LED for providing photosynthetically active radiation to leaf. |
| LED panels | Custom in-house fabrication with basic circuitry (Fig. 11) | Provide photosynthetically active radiation to the leaf |
| LED output control | Custom in-house fabrication with basic electrical control circuitry | Sets and ensures constancy of output of LEDs |
| Heater | Custom in-house fabrication with standard peltier device and thermostat (Fig. 4) | Regulates instrument and sample gas temperature |
| Cuvettes | Custom designed by the authors; fabricated by Tucson Store Fixtures, Tucson, AZ, USA (Fig. 8) | Flow-through acrylic cuvettes enclose leaves to capture emitted gases |
| Plastic cases | Fuerte Cases, El Cajon, CA, USA | "Seahorse" waterproof plastic cases, used to house PORCO components. |
| Dehumidifier | Custom in-house fabrication, see Fig. S1 | Reduces sample gas humidity to low and constant level |

**Table S2:** (Dataset not provided with bioRxiv pre-print.) This separate file contains data representing the capacity to emit isoprene or not, among tropical tree species. Data from the present study is reported, as well as published data from tropical species that share a genus with those reported in the present study. The previously published data is the source of the calculated proportion of species per genus that emit isoprene, reported in Table 1 of the present study. The published data is derived from a combination of gas-distinguishing and non-distinguishing detection methods. Substantial light-dependent emissions detected by a non-distinguishing method are assumed to be isoprene—the most abundantly emitted volatile isoprenoid. Species measured in the present study are reported as isoprene 'emitters' where emissions were >1 nmol $m^{-2}$ $s^{-1}$. Taxonomic names from all sources were standardized using the Taxonomic Name Resolution Service (tnrs.iplantcollaborative.org). Specimen collection numbers from the present study are provided, and herbarium registration IDs for voucher specimens where available (fertile specimens only). Non-fertile specimens are stored at a separate teaching herbarium. Contact the corresponding author for questions about species identifications and non-fertile voucher specimen accessibility. Detailed specimen collection notes and photos are available upon request. All data processing was performed with code in R, including detailed quality control, so data-handling errors should be minimal.

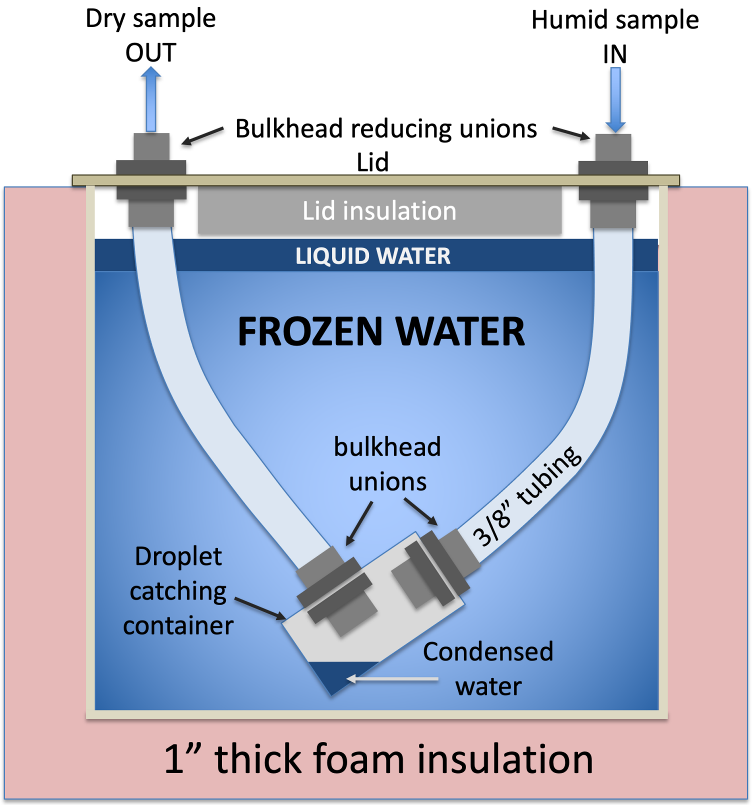


**Figure S1:** Cartoon of sample-gas dehumidifier used in PORCO, capable of maintaining low and constant relative humidity (RH; approximately 10 +/- 0.5 % RH at the PID inlet) over 16 hours of continuous run time in hot (30-40˚C air) conditions. The dehumidifier is labeled as Item 6 in Fig. 1. This system was optimized over many iterations, and requires particular features and procedures to be successful. Water vapor is condensed out of sample air by routing air through tubing embedded in ice and water. The container is prepared by filling with 3-4 L of water, submerging the tubing assembly attached to the lid, and freezing for 36 hrs. With appropriate gaskets and careful seals, the freezing process does not cause the assembly to leak, and PTFE or PFA tubing will not crack. Achieving constant RH requires that the temperature of the assembly be held at exactly 0 ˚C by the thermal equilibrium of ice and water. Therefore, before starting measurements, it is critical to partially melt the ice by running water over the outside of the container, and adding some liquid water to the top. Otherwise, the ice temperature will gradually rise from a starting temperature < 0 ˚C, changing the saturated vapor pressure, until an ice temperature of 0 ˚C is reached. Tubing diameter smaller than 3/8" results in water vapor freezing against the tubing walls and becoming ice plugs that block air flow. The path length is approximately 14". Larger diameters and longer path lengths result in larger mixing volumes that temporally attenuate the gas signal. A small droplet-catching chamber captures condensed water, allowing the sample gas to pass above it. The chamber is emptied after each day of measurements. As soon as some ambient vapor is condensed in the droplet catcher, the liquid water will humidify dry calibration gas so that calibration RH matches sample RH (see discussion of 'PID drift').

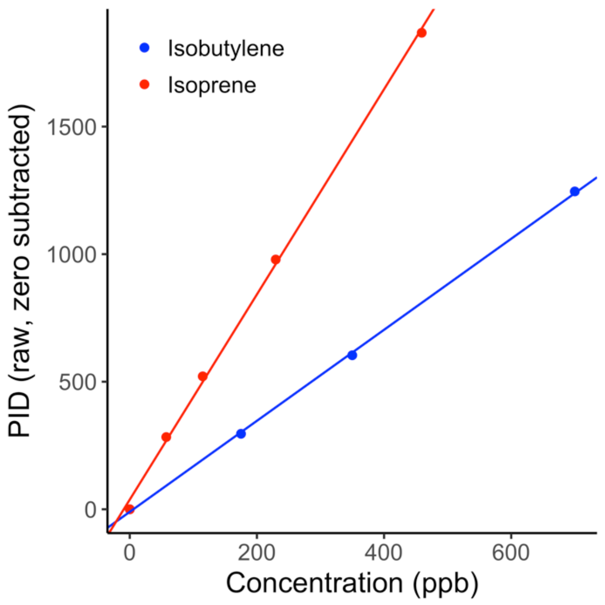


**Figure S2:** PID response to isoprene standard (red, slope = 4.020) compared to isobutylene standard (blue, slope = 1.784). Relative sensitivity to the two gases requires a correction factor of 0.444 when measuring isoprene following isobutylene calibration.

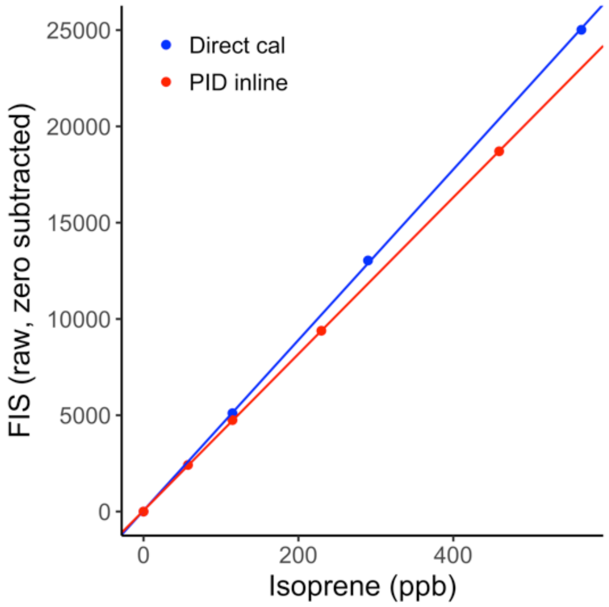


**Figure S3:** Isoprene calibration of FIS by direct application (blue, slope = 44.28) or with PID inline (red, slope = 40.65). The PID reduces isoprene concentrations by 8.2%.

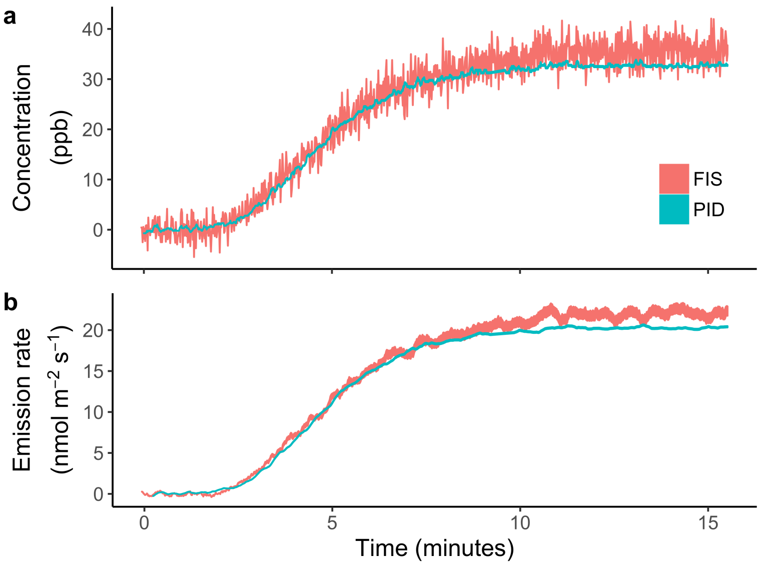


**Figure S4:** Isoprene emission measured from a leaf of *Malpighia glabra* by Fast Isoprene Sensor (FIS) and photoionization detector (PID) incorporated in the PORCO system. Emissions are sampled from a Licor-6400 leaf cuvette, drawn first through the PID and then through the FIS. Panel (a) shows concentration data from both instruments, demonstrating greater signal noise in the FIS compared to the PID. Panel (b) shows emission rates calculated from the concentration data.

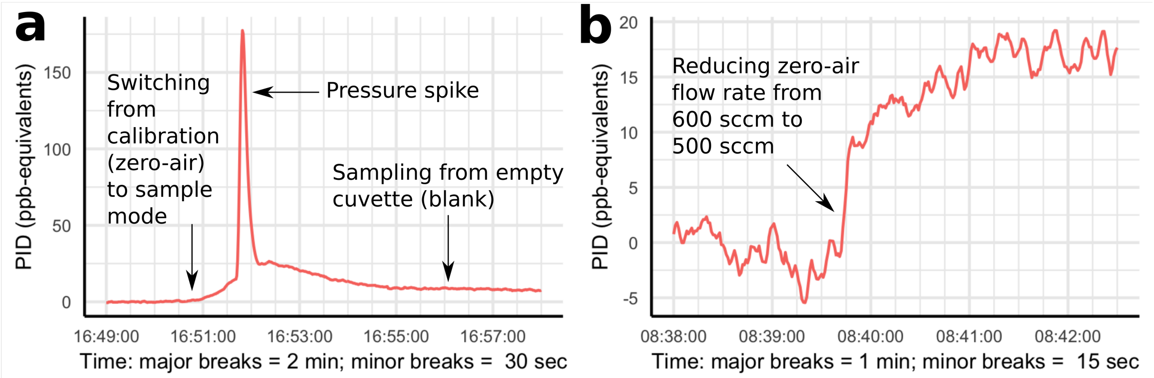


**Figure S5:** Examples of PID sensitivity to sample air pressure. PID signal is represented in calibrated ppb-equivalents, though only purified 'zero air' is measured in both panels. Shown in (a), a "pressure spike" in PID readings occurs when changing valves to alter flow paths between calibration and sample modes, or any time the PID air pump meets resistance due to blockage of a flow path. Shown is a moderate and transient spike, but more extreme pressure changes can cause spikes equivalent to >1 ppm and require more than a half hour to recover. Shown in (b), a small change in positive pressure due to changing the zero-air flow rate in calibration mode causes a step-change in baseline readings. The diagram of flow configurations in Fig. 1 (Main Text) shows why the PID experiences positive pressure in calibration mode (air is forced from a pump and compressed calibration gas bottle) and negative pressure in sample mode (PID onboard pump pulls air from cuvette through dehumidifier and thermal equilibration tubing).

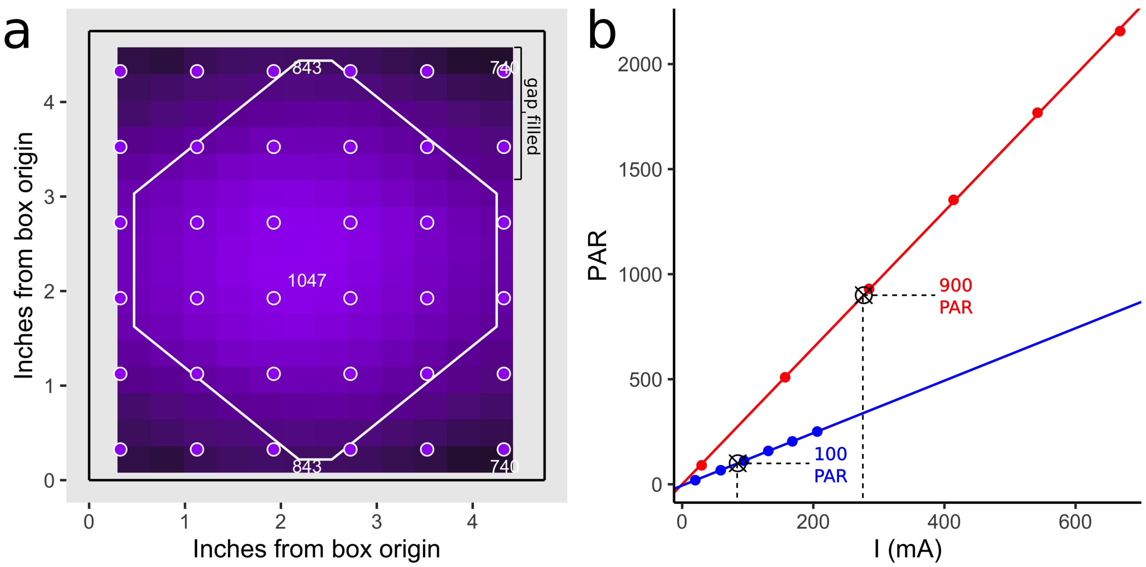


**Figure S6:** The light emitting diode (LED) light source was calibrated to provide a precise and even light distribution over the leaf plane. (a) Light was measured at the leaf plane using a photosynthetically active radiation (PAR) sensor on a measurement grid inside of a tall, square cuvette. The custom LED panels were mounted atop the cuvette, above a diffusion plate as under field conditions. The target leaf region is outlined in white. (b) A calibration between PAR and electrical current (I) for each color of LED in the central region of the measurement grid allows precise control over light output. By relating light intensity in the central (calibration) region to average light intensity over the leaf region (white polygon), we determined the electrical current required to produce an average of 1000 PAR at the leaf plane, in proportions of 90% red light and 10% blue light (matching the LICOR-6400 photosynthesis chamber light output).

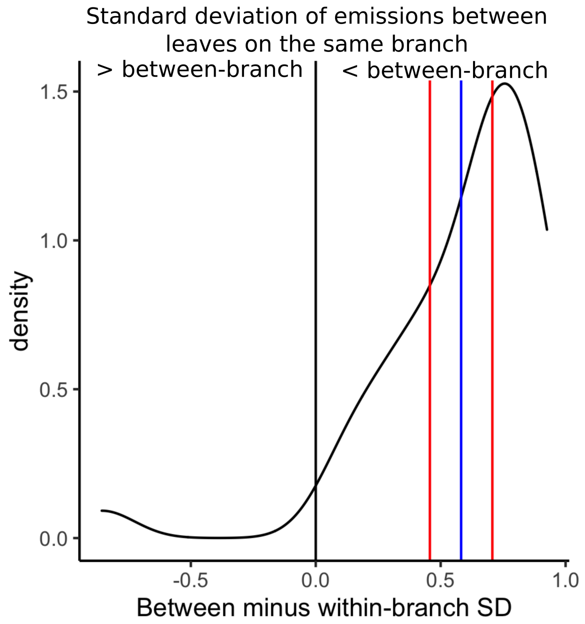


**Figure S7:** Between-branch variation in emission rates (SD = 0.926) exceeds within-branch variation (mean SD = 0.58) by a factor of 1.6 (t-test, $p << 0.001$). Leaf emission rates (nmol $m^{-2}$ $s^{-1}$ +1) were log transformed after converting emission rates below 0.4 to 0 (below detection limit), and aggregated to the branch scale. Branch data is filtered to species with a maximum emission rate ≥ 1. The distribution shown is the standard deviation of branch-mean emission rates (i.e. between-branch variation) minus the standard deviations of leaf emissions per branch (i.e. within-branch variation). The distribution significantly exceeds zero, with blue and red lines showing the mean and 95 % CI estimated by t-test. This demonstrates that PORCO is capable of differentiating ecologically driven variation in emission rates which manifest at the branch scale, against stochastic variation in individual leaf emission rates and measurement error.
